# Deep learning with implicit handling of tissue-specific phenomena predicts tumor DNA accessibility and immune activity

**DOI:** 10.1101/229385

**Authors:** Kamil Wnuk, Jeremi Sudol, Kevin B. Givechian, Patrick Soon-Shiong, Shahrooz Rabizadeh, Christopher Szeto, Charles Vaske

## Abstract

DNA accessibility is a key dynamic feature of chromatin regulation that can potentiate transcriptional events and tumor progression. Recently, neural networks have begun to make it possible to explore the impact of mutations on DNA accessibility and transcriptional regulation by demonstrating state-of-the-art prediction of chromatin features from DNA sequence data in specific tissue types. We demonstrate enhancements to improve such tissue-specific prediction performance, and show that by extending models with RNA-seq expression input, they can be applied to novel tissue samples whose types were not present in training. We show that our expression-informed model achieved particularly consistent accuracy predicting DNA accessibility at promoter and promoter flank regions of the genome.

Leveraging this new tool to analyze tumor genomes across tissues, we provide a first glimpse of the DNA accessibility landscape across The Cancer Genome Atlas (TCGA). Our analysis of the Lung Adenocarcinoma (LUAD) cohort reveals that viewing tumors from the perspective of accessibility at promoters uniquely highlights several immune pathways inversely correlated with an overall more open chromatin state. Further, through identification of accessibility sites linked with differential gene expression in immune-inflamed LUAD tumors and training of a classifier ensemble, we show that patterns of predicted chromatin state are discriminative of immune activity across many tumor types, with direct implications for patient prognosis. We see such models playing a significant future role in matching patients to appropriate immunotherapy treatment regimens, as well as in analysis of other conditions where epigenetic state may play a significant role.

**Significance Statement:** DNA accessibility determines whether proteins have access to DNA-binding sites and is a key dynamic feature that influences regulation of gene expression that differentiates cells. We improve and extend a neural network model in a way that expands its application domain beyond studying the impact of genetic sequence and mutations on DNA accessibility in specific cell types, to tissues for which training data is unavailable.

Leveraging our tool to analyze tumor genomes, we demonstrate that in lung adenocarcinomas the accessibility perspective uniquely highlights immune pathways inversely correlated with a more accessible DNA state. Further, we show that accessibility patterns learned from even a single tumor type can discriminate immune inflammation across many cancers, often with direct relation to patient prognosis.

## Introduction

DNA accessibility plays a key role in the regulatory machinery of DNA transcription. Locations where DNA is not tightly bound in nucleosomes - detectable as DNase I hypersensitivity sites (DHSs) - render the sequence accessible to other DNA-binding proteins, including a wide range of transcription factors (TFs). DHSs are cell-specific and play a crucial role in determining transcriptional events that differentiate cells.

Furthermore, genome-wide association studies (GWAS) have revealed that the vast majority of genetic variants significantly associated with many diseases and traits are located in non-coding regions (2) and well over half non-coding single nucleotide polymorphisms (SNPs) affect DHSs (3). Thus variable access to DNA regulatory elements not only plays a key role in normal cell development, but also in altered expression profiles associated with disease states (2, 4), including cancer.

In an effort to go beyond association studies and gain deeper insight into how changes in DNA sequence impact transcriptional regulation, some groups have developed predictive models for a multitude of genomic phenomena. Several works have recently made significant advances in accuracy of such DNA-sequence-based prediction tasks by applying neural network models to problems such as transcription factor binding (5–8), promoter-enhancer interactions (9), DNA accessibility (6, 10, 11), and DNA methylation states (11, 12).

One common issue that limits the broad applicability of these models is the cell-type-specific nature of many of the underlying biological mechanisms, such as DHSs. All the above examples approached their prediction problems by learning to estimate the conditional probability *p*(*a*|***d***, *b*), where *a* is the accessibility (or other attributes) of a segment of DNA sequence, ***d***, and *b* is a discrete label of tissue type. In practice this meant either training a separate model for each cell or tissue type, or having a single model output multiple tissue-specific (multi-task) predictions. This made it difficult to apply the models to new data and limits them from being integrated into broader scope pathway models (13).

Conveniently, a number of studies have demonstrated that gene expression levels from RNA-seq can be used to discriminate cell types (14–16), providing evidence that *p*(*b*|***r***) can be learned (where ***r*** is a vector of RNA-seq gene expression measurements). In addition, DNase-seq and microarray-based gene expression levels from matched samples were found to cluster similarly according to biological relationships, with many DHSs found to significantly correlate with gene expressions (17).

Our work focuses on overcoming the barrier to broad applicability due to cell-type-specific phenomena by putting the burden on a deep neural network classifier to handle the complex relationship between expression and DNA sequence accessibility without intermediate discrete tissue labels. Our model directly estimates *p*(*a*|***d***, ***r***), and thus implicitly handles the space of possible tissue types and states, *B*, since *p*(*a*|***d***, ***r***) = Σ_*i*∈*B*_ *p*(*a*|***d***, *b*_*i*_)*p*(*b*_*i*_|***r***). This allows the model to exploit similarity information in the space of tissue types and make predictions for previously unseen tissues whose gene expressions were similar but unique from samples in the training data.

We build on the Basset neural network architecture of Kelley *et al* which recently demonstrated state-of-the-art results on DNA accessibility prediction (10). They factored the cell-specific DHS issue into their work by first creating a binary matrix of sample tissues and their respective accessibility for a list of genomic sites. The universal list of (potentially accessible) sites was found by agglomerative clustering all overlapping DNase-seq peaks across all samples before training. These potentially accessible sites defined the 600 base-pair DNA segments used as inputs. Second, they set up the model’s final layer as a multi-task output, with a distinct prediction unit for each tissue type.

We began by showing that neural network performance on the original task of Kelley *et al*, predicting accessibility at held out sites across 164 tissue types, could be improved by strategic factorization of convolutional layers. Then, based on our hypothesis that a neural network should be capable of learning to appropriately modulate its prediction of the DNase-seq signal if informed with RNA-seq, we extended the network to handle a vector of gene expression values as input. To train this model we constructed a new dataset of samples from the ENCODE project (18) consisting only of RNA-seq and DNase-seq measurements whose correspondence could be determined. Our new model achieved compelling results for held-out tissue types, and proved to be very reliable at predicting accessibility at promoter and promoter flank genomic regions.

We then applied our accessibility prediction model to whole genomes across six cancer cohorts from The Cancer Genome Atlas (TCGA) (19), as summarized in Figure 1A, and highlighted that accessibility complements RNA-seq. For example, clustering lung adenocarcinoma (LUAD) samples based on predicted accessibility was distinct from any RNA-seq-based cluster assignment, and revealed a group of patients showing enrichment for pathways involved in immune response. Further, splitting the same cohort by immune cell composition revealed a difference in survival and enabled training of a classifier to detect an immune-inflamed tumor state in LUAD, based only on accessibility predictions for a small set of sites. This classifier allowed us to discriminate immune active patient groups with significant differences in survival in several distinct cancers, often aligning with findings from other cancer immunology studies. To the extent of our knowledge, this is the first time that a prediction of DNA accessibility has been applied to whole genomes in TCGA cancer cohorts to infer the chromatin landscape across cancers.

**Figure 1.**
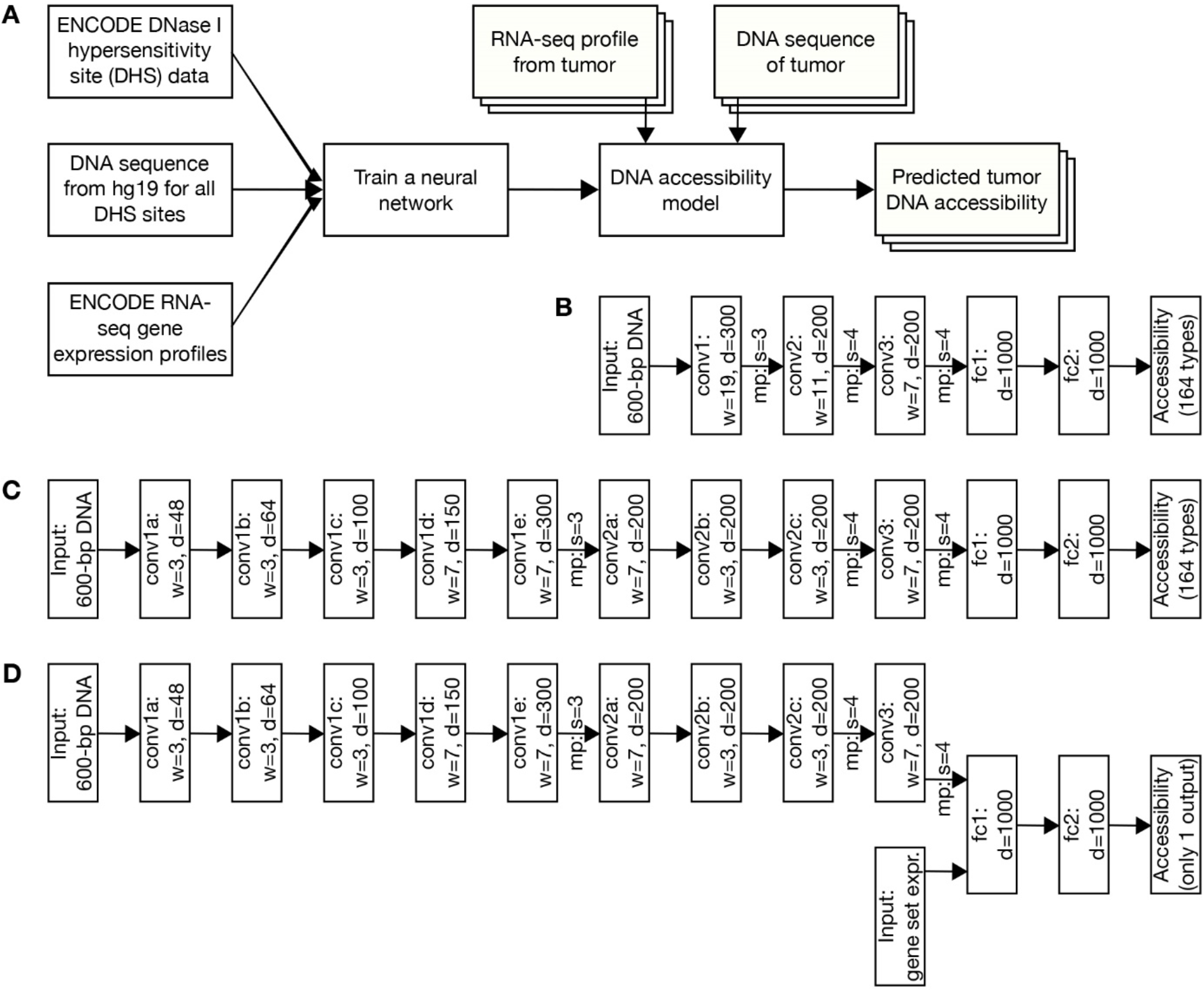
Overview of our pipeline from training to application. (*A*) DHS, hg19 DNA and RNA-seq information are all used to train the neural network. With tumor RNA-seq and DNA sequence data input the DNA accessibility model can then be used to predict chromatin state in tumors. (*B*) The neural network architectures for the tissue-specific baseline model, (*C*) the tissue-specific factorized convolutions model, and (*D*) the expression-informed model are shown. Depth (d =) is provided for all fully connected (fc) layers. Convolution (conv) layers also list their width (w =). Max pooling (mp) is indicated where present between layers and is always applied with equal size and stride (s =).

We anticipate our expression-informed model may not only provide detailed information regarding DNA accessibility across tissues and enable discrimination of immune-inflamed tumors, but might also be used to predict individual patient response to various immune-based therapies. We also expect our approach to be a useful tool in understanding of other conditions where chromatin state is suspected to play an important role including aging (20), neurodegenerative disease (21), autoimmune diseases (22), as well as autism (23). Finally, we stress that our approach to implicitly handling tissue type and state can be used in any DNA-based prediction task.

## Results

### Convolutional layer factorization improves accuracy

As a baseline we first implemented our own version of the Basset architecture (Figure 1B), with minor changes not related to network structure. Compared to an ROC AUC = 0.895 reported by Kelley *et al* on their benchmark test set (10), and confirmed by applying the pre-trained model provided by the authors, our baseline implementation achieved a mean ROC AUC = 0.903 (Table 1).

**Table 1.** Tissue-specific model results on Basset benchmark dataset

| Tissue-specific model | mean ROC AUC | mean PR AUC |
| --- | --- | --- |
| Basset (pre-trained) | 0.895 | 0.561 |
| Our baseline | 0.903 | 0.582 |
| Our factorized convolutions | 0.910 | 0.605 |

Following the success of many works demonstrating that deeper hierarchies of small convolutional kernels tend to improve neural network performance (24–26), we experimented with factorization of large convolutional layers in the baseline model. We found that factorization of layers closest to the data was the most significant for improving accuracy (Supplemental Figure S1). When only the second convolutional layer was factorized, the speed of learning improved during the early epochs of training, but final accuracy was not noticeably affected compared to our baseline implementation. An overall improvement in both rate of learning and final accuracy was achieved when both the first and second convolutional layers were factorized (Figure 1C), leading to a mean ROC AUC = 0.910 (Table 1).

Furthermore, despite following the same training procedure and taking no additional steps to account for class imbalance, our final tissue-specific model’s mean PR AUC = 0.605 compared favorably to the mean PR AUC = 0.561 reported as the best result obtained by the Basset model.

### ENCODE DNase-seq and RNA-seq dataset

In order to train a model for predicting accessibility that is implicitly informed about tissue state through gene expression it was necessary to build a new dataset where both DNase-seq and RNA-seq were available for a large and diverse collection of different tissue types. We collected all human samples from the ENCODE project (18) for which correspondence between RNA-seq and DNase-seq measurements could be determined. After errors were filtered out, the final dataset consisted of 74 unique tissue types, with 220 DNase-seq files and 304 RNA-seq files. A validation set of 5% randomly held out samples was split from the data so that tissue types were diverse but still independent measurements of tissues also appearing in training.

Two sets of paired test and training partitions were created (Supplemental Figure S2). The first partition pair (tissue overlap in Table 2) was constructed in the same way as the validation data by randomly holding out test samples and allowing for tissue type overlap with training. The second partition pair (held-out tissue in Table 2) was constructed such that the test set was comprised only of samples from tissue types that were not present in either training or validation partitions. The latter was meant to more accurately simulate the intended application scenario and was thus the main focus throughout our analysis.

**Table 2.** File and tissue type distribution per dataset partition

| Partition | Unique tissues | DNase-seq files | RNA-seq files |
| --- | --- | --- | --- |
| Validation | 10 | 11 | 12 |
| Tissue overlap train | 73 | 195 | 277 |
| Tissue overlap test | 12 | 14 | 15 |
| Held-out tissue train | 66 | 198 | 281 |
| Held-out tissue test | 8 | 11 | 11 |

The data partitions were revised once from their first iteration when several erroneous samples were discovered and revoked by the ENCODE consortium. Once revoked samples were removed we saw a significant decrease in spurious DHSs. Table 2 shows the distributions of the final dataset.

### Expression-informed model can predict accessibility in held out tissues

We explored several alternative versions of our expression informed neural network along with a range of different hyperparameters. Based on validation performance, we found that concatenating gene expression directly with output from the convolutional layers, using a large batch size with appropriately matched learning rate, and weight initialization from a model trained using more noisy training data made the most impact. Changing the fraction of positive samples per training batch from 0.5 to 0.25 also led to a minor improvement.

Supplemental Table S2 shows that final model (Figure 1D) performance on the validation set, both overall and by tissue type, was consistent across each of the two training partitions with respect to both ROC AUC as well as PR AUC. Table 3 and Table 4 summarize the results of applying our model across whole genomes, at all potential DHSs. For tissue types with more than a single file pair in the test set, each sample’s results are listed.

**Table 3.** Tissue overlap whole genome test results

| Sample tissue type | ROC AUC | PR AUC |
| --- | --- | --- |
| A172 | 0.959 | 0.721 |
| left renal pelvis | 0.967 | 0.838 |
| small intestine | 0.951, 0.926 | 0.737, 0.571 |
| muscle of arm | 0.968 | 0.843 |
| forelimb muscle | 0.968 | 0.808 |
| keratinocyte | 0.939 | 0.644 |
| skin fibroblast | 0.948, 0.947 | 0.770, 0.763 |
| large intestine | 0.964 | 0.727 |
| muscle of back | 0.967, 0.954 | 0.853, 0.854 |
| adrenal gland | 0.957 | 0.743 |
| SK-N-DZ | 0.898 | 0.652 |
| fibroblast of lung | 0.942 | 0.840 |
| mean tissue type AUC | 0.950 | 0.758 |
| **overall AUC** | **0.947** | **0.748** |

**Table 4.** Held-out tissue whole genome test results

| Sample tissue type | ROC AUC | PR AUC |
| --- | --- | --- |
| left kidney | 0.965 | 0.778 |
| OCI-LY7 | 0.899, 0.899, 0.886, 0.886 | 0.654, 0.654, 0.655, 0.654 |
| prostate gland | 0.865 | 0.516 |
| hindlimb muscle | 0.943 | 0.824 |
| spleen | 0.913 | 0.582 |
| astrocyte | 0.919, 0.944 | 0.787, 0.613 |
| fibroblast of skin of abdomen | 0.964 | 0.826 |
| G401 | 0.739, 0.846 | 0.459, 0.516 |
| mean tissue type AUC | 0.898 | 0.655 |
| **overall AUC** | **0.897** | **0.621** |

As expected, overall the model was less accurate on completely new tissue types; however, even in the more challenging scenario, the overall PR AUC was higher than the best tissue-specific models evaluated on known tissue types. Note that several of the results in Table 4 were within similar ranges as predictions whose sample types overlapped with training.

### Expression-informed model predictions are highly reliable at promoter and promoter flank genomic sites

To better understand the performance characteristics and limitations of our model, we broke down our ENCODE validation and test results by genomic site type. Table 5 details the distribution of annotations applied to the 1.71 million sites considered in the held-out tissue training set, the percent of all positive samples that fall within each annotation, and the percentage of samples per each annotation type that are positive. Note that a single site may overlap with more than one annotation.

**Table 5.** Distribution of potentially accessible sites by annotation

| Site type | % of all sites | % of all positive examples | % per site type that are positive |
| --- | --- | --- | --- |
| exon | 3.47 | 5.08 | 9.74 |
| intragenic | 49.94 | 45.25 | 6.04 |
| intergenic | 39.89 | 34.11 | 5.70 |
| promoter and flank | 6.37 | 30.33 | 31.75 |
| enhancer | 1.08 | 3.81 | 23.47 |
| other | 5.39 | 5.20 | 6.43 |

We found that even for samples in which the model performed poorly overall, predictions within promoter and promoter flank regions consistently attained a high level of accuracy (Figure 2A and D, Table 6), achieving a PR AUC = 0.839 over all held out tissue types and a PR AUC = 0.911 over randomly held out samples (validation set).

**Figure 2.**
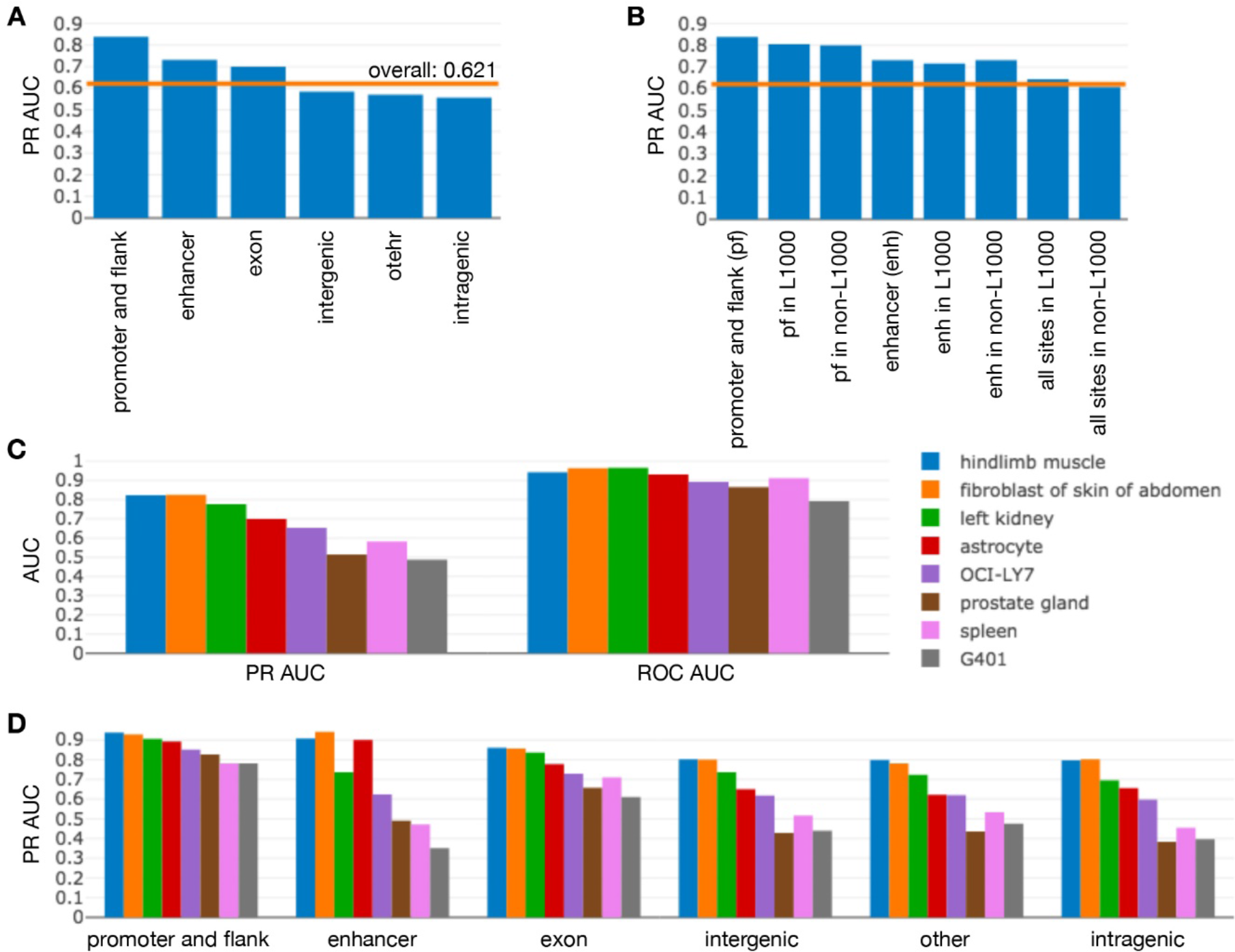
Promoter and promoter flank accessibility (pf) is highly predictable, but enhancers show variability. (*A*) pf accessibility is highly predictable (PR AUC = 0.839), as shown by the genomic site performance breakdown over all samples in the held-out tissues test set. The orange line indicates overall PR AUC computed across all test samples and all sites. (*B*) No clear performance difference was observed when genomic sites across the held-out tissue test set were split into those that did (in L1000) and did not (non-L1000) overlap the L1000 RNA-seq input gene set. Note that not all sites overlapped with known gene regions, so the union of the L1000 and non-L1000 subsets did not always make up the complete set of sites of a certain type. (*C*) Overall metrics separated by tissue type show that some held-out tissues in the test set were more challenging as reflected by lower AUCs. (*D*) Predictions at enhancers were highly variable between samples – even with good PR AUC - and performance on pf regions remained consistently high, even for tissues where overall results were lowest.

**Table 6.** Promoter and promoter flank results across held-out tissue test set whole genomes

| Sample tissue type | ROC AUC | PR AUC |
| --- | --- | --- |
| left kidney | 0.949 | 0.905 |
| OCI-LY7 | 0.869, 0.868, 0.864, 0.864 | 0.842, 0.842, 0.859, 0.859 |
| prostate gland | 0.897 | 0.826 |
| hindlimb muscle | 0.935 | 0.938 |
| spleen | 0.867 | 0.782 |
| astrocyte | 0.925, 0.914 | 0.946, 0.838 |
| fibroblast of skin of abdomen | 0.951 | 0.929 |
| G401 | 0.798, 0.828 | 0.807, 0.757 |
| **overall AUC** | **0.876** | **0.839** |

We also confirmed that the accuracy of these predictions was independent of whether the promoter and promoter flank sites overlapped with the regions of genes used in our RNA-seq input gene set (Figure 2B).

Selecting a threshold for classification of only promoter and promoter flank sites such that precision is 80% (20% false discovery rate) on the held out tissue test set, our trained model recalls 65.3% of accessible promoter regions, with a false positive rate of 10%. Applying this same threshold to the validation set where tissues are allowed to overlap with the training set, the model achieves a precision of 93.4%, recalling 62.6% of accessible promoter regions, and has a false positive rate of only 3.5%.

We also investigated accuracy at enhancer sites, finding a PR AUC = 0.732 over held-out tissues and PR AUC = 0.889 over randomly held out samples (validation set). Differently from promoter and promoter flank regions however, enhancer prediction accuracies showed a high variance between test samples (Figure 2D, Supplemental Table S3). Thus, more investigation is necessary before relying on accessibility predictions at enhancers in further analysis.

To quantify the effect of similarity to training data on prediction performance we looked at correlation between PR AUC (computed independently for all predictions in each whole genome sample) and distance of each test and validation sample to its closest sample in the training set (Table 7). As might be expected from Figure 2D, we confirm that prediction performance is less correlated with test sample similarity to training data at promoter and promoter flank sites than when PR AUC is evaluated over all potentially accessible sites.

**Table 7.**
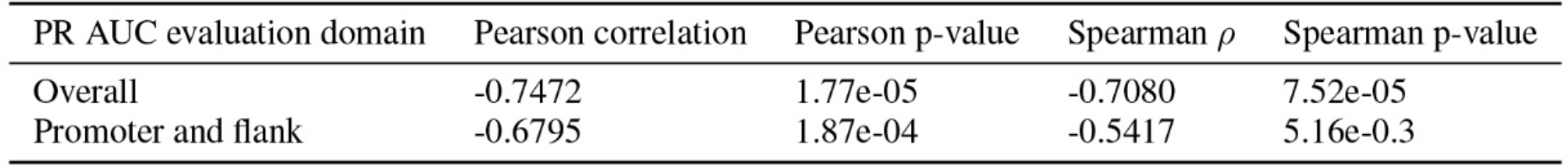
Correlation of PR AUC with test sample distance to the nearest training sample

### Promoter accessibility patterns across cohorts from The Cancer Genome Atlas

We applied our trained model to promoter and promoter flank sites in TCGA samples from six cohorts (LUAD, LUSC, KICH, KIRC, KIRP, and BRCA). Across all samples in these cohorts for which whole genome sequencing (WGS) was available, 3172 interest regions had one single nucleotide polymorphism (SNP), 78 had 2 SNPs, and only 9 regions had between 3 and 5 SNPs. A total of 465 sites included insertion or deletions (INDELs), and only 7 sites featured both an INDEL as well as an SNP. Lung cancers exhibited the highest average number of mutated sites per patient from our selected cohorts (Figure 3A).

**Figure 3.**
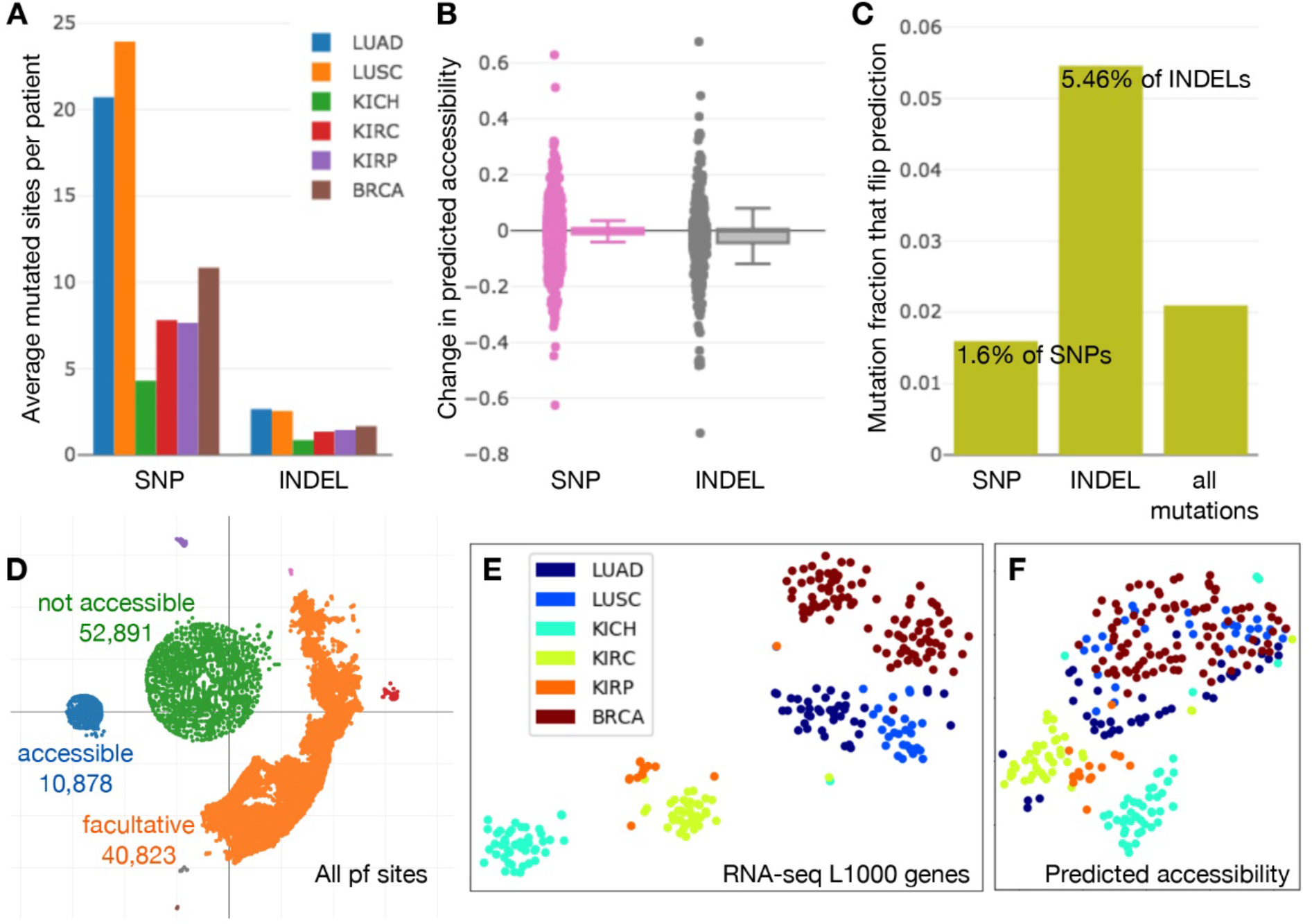
SNP and INDEL mutations and predicted accessibility landscape in tumors. (*A*) The average number of single nucleotide polymorphism (SNP) and insertion or deletion (INDEL) mutations that overlap prediction sites per patient across 6 TCGA cohorts is shown. (*B*) When predictions at sites with mutations were compared with and without applying mutations to the input DNA sequence, the change in predicted accessibility exhibited a higher variance for INDELs than SNPs. (*C*) In addition, a larger fraction of sites with INDELs were responsible for a change in the classification decision (flipped prediction) than for sites with SNPs. (*D*) Using t-SNE (perplexity = 50) to visualize the predicted accessibility of individual pf sites across our selected TCGA samples, we identified which sites were facultative (orange), constitutively accessible (blue), and constitutively not accessible (green). (*E*) Finally, t-SNE applied to patient samples exhibited different relationships (such as a clear split in BRCA samples) when based on RNA-seq gene expression of the L1000 gene set, than (*F*) when based on predicted accessibility at all pf sites within each sample (in which case lung and breast cancers appeared to share some common characteristics).

To observe the effect of region changes on accessibility, we compared predictions with and without SNPs and INDELs present. INDELs had the greatest impact on predicted accessibility, exhibiting a higher variance than SNPs (Figure 3B) and leading to a change in accessibility classification in 5.46% of cases (at the previously defined accessibility threshold that achieved 80% precision) (Figure 3C).

To get a landscape view of how promoter and promoter flank sites behave, we embedded their binary accessibility decisions across all our selected TCGA samples in two dimensions with t-SNE (27). This clearly separated sites into constitutively accessible, constitutively not accessible, and facultative (Figure 3D). A few very small clusters of several hundred sites were also formed, which were groups of typically constitutive sites that acted uniquely in one or two individual patients.

Second, we stacked all predictions into a single vector per patient to form accessibility profiles for all samples in our six TCGA cohorts, and again applied t-SNE to visualize relationships (Figure 3F). This qualitatively showed that looking at cancers from the viewpoint of DNA accessibility highlights different relationships than analysis of RNA-seq alone. For example, in the RNA-seq t-SNE space (Figure 3E) a clear separation emerges among breast cancers (BRCA), which correspond to basal-like versus luminal A/B and HER2-enriched clusters. In contrast, in accessibility t-SNE space (Figure 3F), the lung (LUAD, LUSC) and breast (BRCA) cancer samples appear to share some common characteristics.

One of many potential biological factors that may contribute to overlap in the accessibility space is the impact of hormone activity on both breast and lung tumors; this activity in turn is epigenetically regulated (28), thus some chromatin state patterns could be shared. There also appears to be a slight partition into left and right groups in how lung cancer samples arrange in the embedding space, with LUAD forming a distinct subset away from the LUSC/BRCA modality within the lung/breast supercluster.

### Accessibility is linked to immune activity in lung adenocarcinoma

We subsequently explored the biological associations of our model’s accessibility predictions by examining transcriptomic data from LUAD samples, as this tumor type has been shown to be of particular interest in chromatin accessibility studies due to the impact on progression (29, 30). Upon clustering LUAD samples for which WGS was available according to their predicted accessibility, clear bifurcation into low accessibility (C0, 21 samples) and high accessibility (C1, 20 samples) samples was observed (Figure 4A). Cluster assignment based on predicted accessibility was distinct from any cluster assignments using the same methodology on RNA-seq directly (Figure 4B).

**Figure 4.**
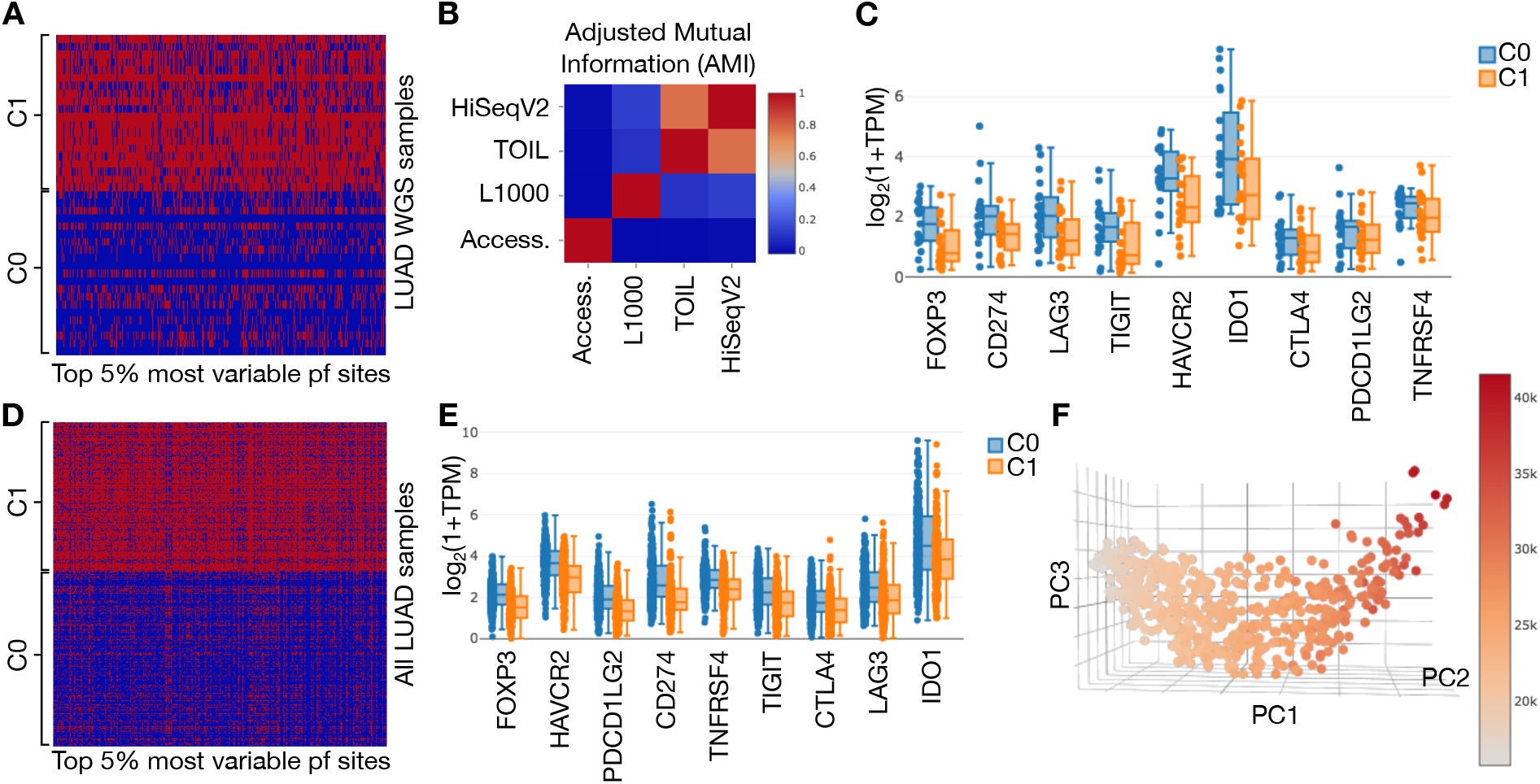
Promoter and promoter flank accessibility and checkpoint gene expression in LUAD WGS samples only and augmented with non-WGS samples. (*A*) The heatmap and patient sample cluster assignment based on the top 5% most variable promoter and promoter flank (pf) accessibility sites across LUAD samples with WGS available are shown. Cluster 0 (C0) has lower overall accessibility (blue = not accessible), and cluster 1 (C1) exhibits generally higher accessibility (red = accessible). (*B*) Adjusted mutual information (AMI) (1) between label assignments based on different data shows higher values (red) between different RNA-seq cluster assignments, and low values (blue) between accessibility (Access.) and clusters based on any other data type. (*C*) Distribution of key checkpoint gene expression levels (with x-axis sorted by significance of two-sided t-test between C0 and C1) shows that the low accessibility group tends to have higher checkpoint levels. (*D*) Applying the same procedure to the full LUAD cohort, which also includes predictions for all non-WGS samples, we see a similar split into low (C0) and high (C1) accessibility groups. (*E*) The same trend in checkpoint expression is observed, with FOXP3 again appearing as the most significant difference (two-sided t-test with Benjamini-Hochberg adjusted p = 4.53e-19). (*F*) Plotting promoter and flank accessibility with respect to its first three principal components (PC1-3) and coloring points by total number of accessible sites in a sample reveals a smoothly varying relationship, motivating a correlation-based approach to exploring the relationship between overall accessibility and gene expression levels.

Differential KEGG pathway expression analysis with Enrichr (31) showed the Chemokine Signaling Pathway (hsa04062) to be upregulated in the low DNA accessibility (C0) patient group. This association held true whether using TOIL RNA-seq data (32, 33) (Enrichr adjusted p-value (adj. p) = 1.191e-6) or HiSeqV2 RNA-seq data [28] (Enrichr adj. p = 0.0145). Chemokines are involved in multiple key processes in tumor growth and immune response (34–36), and their regulation by epigenetic mechanisms has also previously been reported (37, 38).

No difference in tumor mutation burden (TMB) was found between the two clusters (two-sided t-test t = −0.696, p = 0.491), but interestingly the C0 group exhibited higher expression of immune checkpoint genes (Figure 4C). Cell type enrichment analysis (39) of lymphoids and myeloids also revealed a higher level of class-switched memory B-cells (two-sided t-test: t = 4.040, p = 0.000385, Benjamini Hochberg (BH) adj. p = 0.0131) in C0, though estimated levels were generally low in both clusters. Other immune cell estimates exhibited no differences with adj. p < 0.1, which was largely limited by small sample set size.

### Total number of accessible sites correlates with activity in immune pathways

To enhance the scope of our findings, we extended our analysis to all LUAD patient samples for which WGS was not available by predicting accessibility using just the reference genome (hg19/GRCh37) and gene expression data. Although no mutation information was included for these additional samples, only 37 out of 5449 sites (6.79%) used to cluster all WGS data included any instances of mutations. With the additional consideration that only a small percentage of all mutations actually flip binary class predictions (Figure 3C), it is unlikely that cluster assignment of new non-WGS samples was significantly affected by this missing information. As before, the expanded set of patient samples was clustered into two groups according to accessibility.

The group with generally lower accessibility (C0) again exhibited generally higher checkpoint levels (Figure 4D and E). However, visualizing the first three principal components and coloring points by total number of accessible promoter and promoter flank sites (Figure 4F), we did find a smooth change in value along a continuous manifold of samples, primarily along the first principal component (Spearman correlation = 0.989, p = 0.0).

Therefore, instead of differential analysis, all protein-coding genes were filtered by correlation with the total number of accessible sites and evaluated for KEGG pathway enrichment. We found that all genes satisfying the threshold (correlation absolute value > 0.4) had negative correlation values. Since some relationships may not be linear but still monotonic, we focus on the Spearman measure (Supplemental Table S4), although the Pearson measure yielded similar top pathways (Supplemental Table S5). Osteoclast Differentiation (hsa04380, adj. p = 7.45e-15) was the most significantly correlated pathway.

Interestingly, the process of osteoclast differentiation is controlled by two essential cytokines (30): macrophage colony-stimulating factor (M-CSF) and the receptor activator of nuclear factor-κB ligand (RANKL). Tumor Necrosis Factor (TNF) Signaling Pathway (hsa04668, adj. p = 1.30e-8) also appeared among the top three results. TNF has a pro-inflammatory effect and has been noted to play a critical role in the control of apoptosis, angiogenesis, proliferation, invasion, and metastasis (37).

When pathways were sorted by significance, the Chemokine Signaling Pathway - observed in WGS only cluster analysis - appeared eleventh (adj. p = 4.01e-5). Other notable pathways appearing in the top ten included Pathways in Cancer (hsa05200, adj. p = 6.50e-7), Regulation of Actin Cytoskeleton (hsa04810, adj. p = 6.50e-7), NF-kappa B Signaling Pathway (hsa04064, adj. p = 2.68e-7), and Epstein-Barr Virus Infection (hsa05169, adj. p = 2.83e-5). Focal Adhesion (hsa04510, adj. p-value = 2.29e-4) appeared twentieth, but is worth noting as it resurfaces in later analysis.

### Majority of genes with differential accessibility exhibit consistent differential expression in immune cell driven clusters

To investigate accessibility patterns specifically in the context of different tumor immune environments, all LUAD samples were clustered into two groups according to lymphoid and myeloid levels based on xCell cell type enrichment analysis (39). Lymphoid and myeloid cells were selected for their roles in the adaptive and innate immune system, respectively. A thin margin was introduced between clusters to exclude samples with near ambiguous label assignment (Supplemental Figure S6).

Patients in X0 (141 samples) were enriched for many immune cells (Figure 5A), as well as checkpoint gene expression (Figure 5C), and tended to have narrower distributions of both number of accessible promoter and flank sites (generally lower than X1, two sided t-test p = 1.07e-3) and total overall methylation (generally higher than X1, two sided t-test p = 1.29e-7) (Figure 5B). These samples also reflected significantly favorable survival (Figure 5G). We therefore interpreted X0 as the group of “immune hot” patients in LUAD, and X1 as “immune cold”.

**Figure 5.**
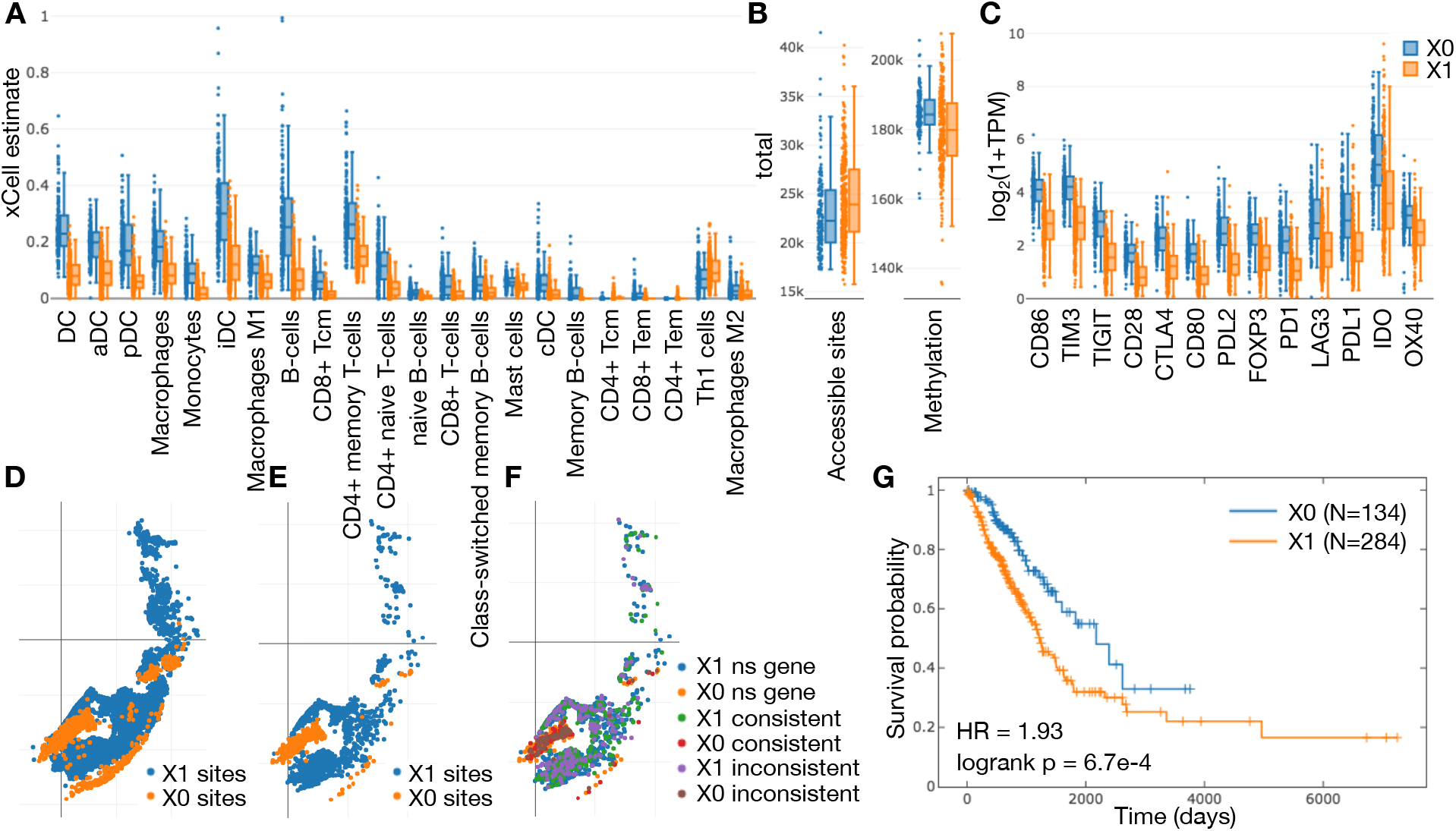
Enrichment in LUAD clusters (after adding a small margin) by cell type, checkpoint expression, methylation, accessibility and survival. (*A*) Cell type enrichment distributions sorted by significance of two-sided t-test for the two clusters (X0, X1) based on xCell lymphoid and myeloid cells with Benjamini-Hochberg adjusted p-value < 1.0e-5 are shown. (*B*) Total number of accessible promoter and promoter flank sites in each sample by cluster (two-sided t-test p = 1.07e-3) along with total methylation (two-sided t-test p = 1.29e-7). (*C*) Checkpoint expression distributions, likewise sorted by significance, also point to a general difference in immune landscape between the two groups. (*D*) All sites with differences in accessibility based on a two-sided t-test with Benjamini-Hochberg adjusted p-values < 0.01 and (*E*) < 1.0e-5 are illustrated on the t-SNE plot of promoter and promoter flank facultative sites. Sites with a difference satisfying the thresholds were assigned to the cluster in which they were more accessible. (*F*) Accessibility differences are further broken down by how they align with direction of upregulation of corresponding nearby genes (ns gene = no significant difference in matching gene, consistent = direction of significant accessibility and gene expression differences are consistent, inconsistent = direction of significant accessibility and gene expression are inconsistent). (*G*) Kaplan-Meier plots demonstrate better survival among X0 (immune hot) patients, shown with logrank test p-value and hazard ratio (HR) based on a Cox proportional hazards (CoxPH) model regression using class assignment as the only explanatory variable.

After eliminating sites that exhibited low standard deviation, we selected all significantly differentiated accessibility sites between the two clusters and mapped them to their nearest gene. Qualitatively we observed that several groups of sites act together in different ways and that those different clusters of chromatin state behavior are stable across significance thresholds (Figure 5D and E). We found that when a majority of sites corresponding to a single gene were accessible more frequently in one cluster, that gene exhibited upregulated expression in the same cluster most of the time (64.7% for genes more accessible in X0 and 64.2% in X1).

Genes whose expression was consistent with increased accessibility in X0 showed near significant levels of enrichment for some pathways that had previously surfaced in our correlation results such as Focal Adhesion (hsa04510, adj. p = 0.0355) and Osteoclast Differentiation (hsa04380, adj. p = 0.0936) (Supplemental Table S6). No significant pathways were found for genes consistent with increased accessibility in X1, or those inconsistent with more accessibility in X0. The strongest significance in pathway enrichment existed in the set of genes inconsistent with increased accessibility in X1 (up in X0 despite accessibility predictions voting for upregulation in X1) (Supplemental Table S7). The most prominent of the enriched pathways in this group were Platelet Activation (hsa04611, adj. p = 4.38e-4), Inflammatory Mediator Regulation of TRP Channels (hsa0475, adj. p = 0.0109), and several with adj. p = 0.0235: Chemokine Signaling Pathway (hsa04062), Focal Adhesion (hsa04510), cGMP-PKG Signaling Pathway (hsa04022), Intestinal Immune Network for IgA Production (hsa04672), Vascular Smooth Muscle Contraction (hsa04270).

These findings suggest that in LUAD tumors, partial regulation of immune- and cytokine-controlled pathways may be exerted via an activator mechanism at promoters. Further, a more significant component of chemokine signaling and platelet activation that distinguishes immune active patients may be subject to repressor regulation at promoter sites.

### Patterns of promoter accessibility predict immune hot tumors with impact on patient survival across several cancers

To further explore the link between DNA accessibility, immune activity, and clinical outcomes we trained an ensemble of three classifiers to detect an immune hot tumor state in LUAD based only on a small subset of accessibility predictions. Applying the ensemble to all of LUAD (Figure 6A) led to a cleaner and more significant partition of patients (compared to Figure 5F and Supplemental Figure S6F) into immune hot and cold tumors. Further applying the classifier ensemble to accessibility predictions across eleven other cancers in TCGA revealed cases where the immune hot state learned from LUAD was beneficial to patient survival (Figure 6 A,B,C,D,E), detrimental to survival (Figure 6 J,K), or had little impact (Figure 6 F,G,H,I,L), with varying degrees of significance.

**Figure 6.**
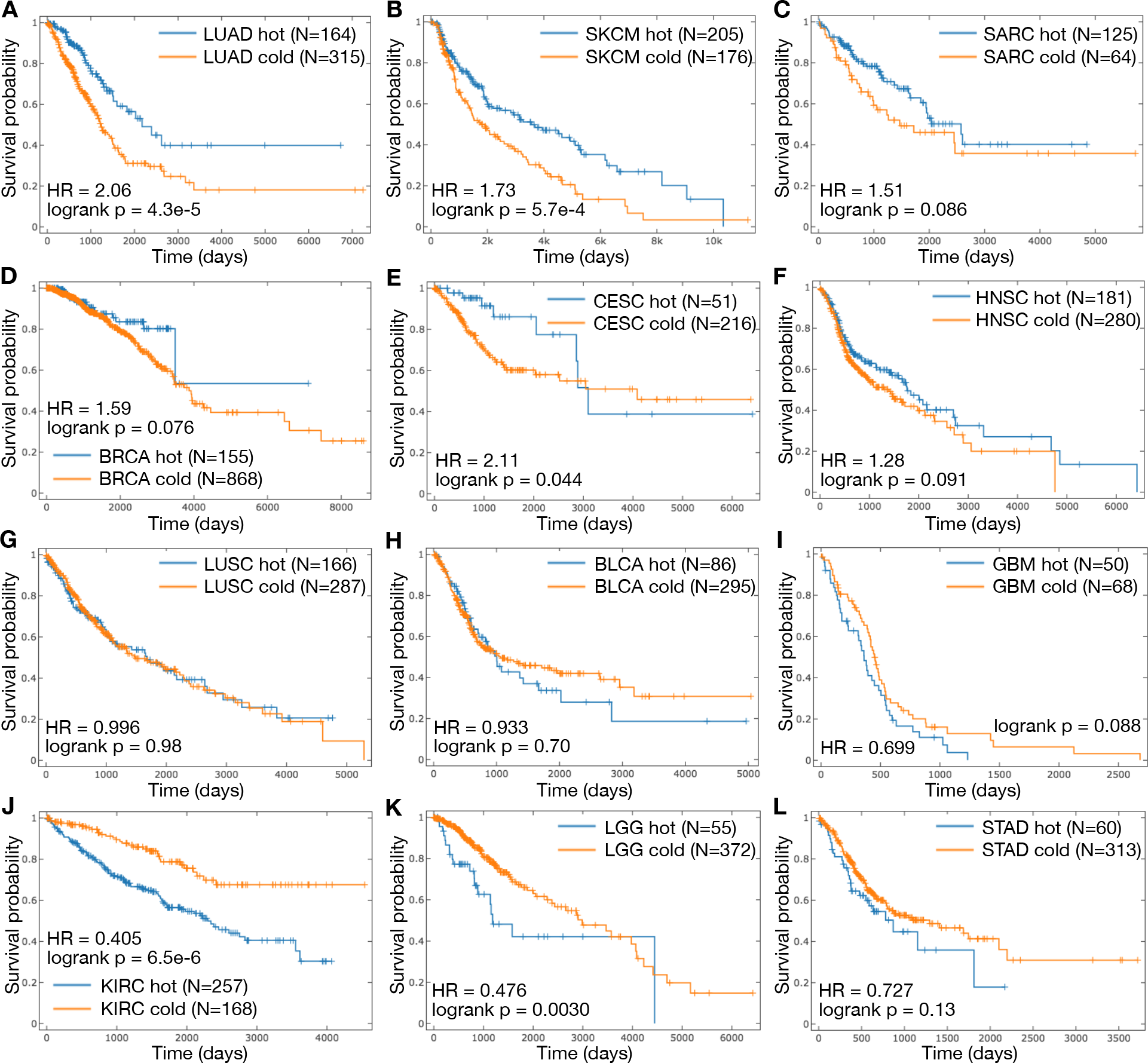
Application of the 3 SVM ensemble for classification of immune hot tumors (trained on subsets of LUAD) with the only input being a vector of 484 promoter and flank predicted accessibility decisions. All Kaplan-Meier plots show group size (N) for patients of both predicted immune activity classes (hot/cold) that satisfy a confidence threshold (see Methods). Also provided are logrank test p-values and hazard ratio (HR) based on a Cox proportional hazards (CoxPH) model regression using class assignment as the only explanatory variable. Note that the time axis range on subplots varies by cohort, and that the immune hot state learned based on LUAD is not always beneficial for patient survival in other tumor types. Tumor types included (*A*) LUAD - lung adenocarcinoma; (*B*) SKCM – skin cutaneous melanoma; (*C*) SARC - sarcoma; (*D*) BRCA – breast invasive carcinoma; (*E*) CESC – cervical squamous cell carcinoma and endocervical adenocarcinoma; (*F*) HNSC – head and neck squamous cell carcinoma; (*G*) LUSC – lung squamous cell carcinoma; (*H*) BLCA – bladder urothelial carcinoma; (*I*) GBM – glioblastoma multiforme; (*J*) KIRC – kidney renal clear cell carcinoma; (*K*) LGG – brain lower grade glioma; and (*L*) STAD – stomach adenocarcinoma.

Our findings aligned very well with a comprehensive analysis of immune subtypes across TCGA (40), which characterized the influence of immune activations on survival. Despite very different methodologies, their plots also indicated that in cohorts such as LUAD, SKCM, and CESC activation of immune subtypes was associated with better outcomes, that the opposite was true in KIRC, LGG, and STAD, and that little impact was visible in LUSC. They did not, however, discuss accessibility as a potential additional biomarker for immune activity.

Significant negative impact of immune activity on survival in KIRC was also shown in a separate cohort of clinical data from Oulu University Hospital (41), confirming that this trend is not unique to TCGA.

Interestingly the study used CD8^+^ T cell count cutoffs to stratify renal cell carcinoma patients into two groups. Based on xCell estimates this was the second most significantly enriched immune cell type (two-sided t-test BH adj. p = 3.57e-26) in KIRC immune hot patients identified by our classifier, after activated dendritic cells (aDC) (two-sided t-test BH adj. p = 1.07e-26) (Supplemental Figure S8). Although the training cohort (LUAD) did express some difference in CD8^+^ T cell enrichment scores between hot and cold tumors (two-sided t-test BH adj. p = 9.65e-15), it was not in the top 10 most significantly different immune cells (Figure 5A). From this we see that our classifiers operating on accessibility predictions learned a more complex decision boundary than simply focusing on direct correlates with the most differentiated immune cell compositions in the training set.

An additional curiosity specific to the KIRC partition from our classifier ensemble is that little difference in CD274 (also called PD-L1) and PDCD1LG2 (also called PD-L2) expression is visible between the two predicted classes, however the strong difference in PDCD1 (also called PD-1) expression levels that exists in other immune hot versus cold partitions does exist. This unique state of checkpoint related gene expression may be linked to the low response rate in renal cell carcinoma patients to anti-PD-L1 therapies, compared to more favorable responses found for anti-PD-1 drugs, in early phase clinical trials (42).

## Discussion

We have demonstrated that predictive models operating on DNA sequence data, extended with RNA-seq expression input, can predict DHSs in unseen tissue types in a way that allows application to new samples without re-training. We showed that these models were capable of achieving consistently high performance for predictions at promoter and promoter flank regions of the genome. Leveraging this new tool for analysis of tumor genomes across different cell and tissue types, we provided the first glimpse of the DNA accessibility landscape across TCGA data.

DNA accessibility is one of many factors that determine expression, which makes inversion of the relationship not trivial; knowing expression levels does not uniquely define the pattern of DHSs. Our expression-informed model (Figure 1D) learns a most likely mechanism by which the DNA sequence immediately surrounding a potential DHS determines its accessibility, conditioned also on an observed expression state. Therefore accessibility prediction applied across the whole genome is an approach to approximately invert gene expression to obtain most likely DHSs.

Our results showed that viewing tumors by promoter accessibility highlights immune pathways that would otherwise be harder to detect from completely unsupervised analysis of RNA-seq data alone. For example, we found several pathways inversely correlated with an overall more open chromatin state. Through identification of facultative accessibility sites linked with differential gene expression in immune-inflamed LUAD tumors and training of a classifier ensemble, we showed that patterns of predicted chromatin state at a small subset of genomic regions are predictive of immune activity across many tumor types, with direct implications for patient prognosis. We see such predictive models playing a significant future role in matching patients to appropriate immunotherapy treatment regimens, as well as in analysis of other conditions where epigenetic state may play a significant role such as autoimmune disease, autism, aging, and neurodegenerative disease.

It may also be interesting to pursue a deeper functional investigation of genes linked with accessibility. Genes with consistent behavior to accessibility are candidates that may be regulated via an activator mechanism at promoters, whereas genes with inconsistent behavior may be subject to alternative gene repression mechanisms, e.g., silencer elements or suppression via microRNAs.

In a few TCGA cohorts, our ensemble classification approach only identified a very small number of immune hot tumor samples, making survival analysis impossible. The generalizability of our immune-related chromatin state across cancers was undoubtedly limited by only having trained the SVMs on a single cohort, since immune cell composition and definition of an immune active state varies across cancers (Thorsson et al. 2018); going forward, we will integrate accessibility signatures from multiple cohorts to train a more comprehensive subtyping of immune state.

Ideally, whole genome sequencing for each of the samples in our training dataset should have been used to learn the most faithful representation of the true biology, as using only reference genome data introduces non-random noise in the input space. Unfortunately, such individual whole genome data was not available for this project. Nonetheless, our work and that of others demonstrates that useful predictors can be learned despite this noise. Unlike models with multi-task outputs, our architecture can easily support such individualized training without any changes and when possible, it will be instituted in the future.

We saw high variance for enhancer sites, but these sites are also interesting with respect to chromatin state and immunotherapy, since they have been linked with T cell dysfunction with potential for therapeutic reprogrammability in mice (Philip et al. 2017). At this time, it needs to be determined whether the large variance in performance is due to limitations in the model, noise in the data, or lack of necessary information in the available inputs. To this end, we look forward to future exploration of a more complete set of genes instead of a manually curated set, such as the LINCS L1000. Many alternatives exist to learn gene embeddings as part of model training, and we believe that ultimately an approach that efficiently incorporates all genes as input will be most effective.

Furthermore, there are a multitude of alternative model architectures such as residual connections (He et al. 2016), densely connected convolutional networks (Huang et al. 2016), and recurrent neural networks (Hochreiter and Schmidhuber 1997) with additions such as attention (Bahdanau et al. 2014; Xu et al. 2015), which we believe are likely to improve performance of our model. These have been left for future evaluation, as the key contribution of this work was movement beyond the cell-type-specific limitations of DNA sequence classifiers, demonstration of the application of our expression-informed model to predict accessibility, and the ability of these predictions to distinguish prognostically alternative immune states across human cancers.

## Methods

### Baseline tissue-specific dataset

All tissue-specific models described in this work were trained and evaluated following the exact procedure of the Basset network (10), using DNase-seq peak data from 164 sample types obtained from ENCODE (18) and Roadmap Epigenomics (43) projects.

Greedy merging of overlapping peaks across all DNase-seq data samples allowed us to create a universal set of potential accessibility sites. For each site, a binary vector was used to label its accessibility state in each of the 164 cell types. Data was then split by genomic site so that 70,000 peak locations were held out for validation, 71,886 for testing, and the remaining 1.8 million sites were used for training. The model input was a 600 base pair window of the DNA sequence centered at a site of interest, represented as one-hot encoding.

### Baseline tissue-specific model implementation

We used TensorFlow (44) to implement the Basset architecture for our baseline. We used Adam (45) instead of RMSProp (46) to optimize network parameters. We also found that use of a dynamic decay rate (that increased over the course of training) for updating moving averages in batch normalization (47) led to a model with competitive performance more quickly than when using a fixed decay. No other significant deviations from the original implementation were included.

When improving on the baseline model with convolutional layer factorizations, we focused experiments on factorizations that maintained the effective region of influence of the original layers and did not significantly increase the overall number of network parameters.

### ENCODE DNase-seq and RNA-seq dataset

Data from the ENCODE project was initially collected at the start of 2017 for all cell or tissue types for which RNA-seq and DNase-seq measurements were both available. In order to capture a greater diversity, gene quantifications from RNA-seq files with the following ENCODE labels were collected: “RNA-seq”, “polyA mRNA”, “polyA depleted”, “single cell”. All files with “ERROR” audit flags were rejected. We kept files with “insufficient read depth,” and “insufficient read length” warnings. Despite being below ENCODE project standards, we believe the available read depths and lengths in warning situations were likely to be less of an issue when it comes to differentiating cell types (16), and preferred to accept more potential noise in favor of a larger diversity of sample types.

The final step of data preparation involved assigning associations between specific RNA-seq and DNase-seq files within the same tissue type. In cases where there existed multiple exact matches of “biosample accession” identifiers between the two file types, associations were restricted to such exact matches. If exact accessions did not match, two file types were associated if it could be verified that they originated from the same tissue sample, cell line, or patient. This eliminated several tissue types for which no such correspondences existed. Both technical and biological replicates were treated as independent samples of the same tissue since we wanted to put the burden of learning non-invertible aspects of noise due to the measurement process on the neural network model.

The dataset was refined in late 2017, as several samples that had been part of our training and testing data were revoked by the ENCODE consortium due to quality concerns and updates. The final dataset consisted of 74 unique tissue types, distributed among partitions as discussed earlier (Table 2). The validation set was held constant, while the training and test sets included two variations.

We utilized the same greedy merge methodology described in Basset (10) on all DNase-seq samples in our training sets to obtain a set of all potential sites of accessible DNA along the whole genome.

We used a fixed length of 600 base pairs (bp) centered at DHS peaks to define each site. Blacklisted sites at which measurements were suggested to be unreliable were excluded (48). This led to a total of 1.71 million sites of interest in the case of the held-out tissue data partition, and 1.75 million sites in the tissue overlap data partition. Using all sites across all available DNase-seq files, this produced a total 338.7 million training examples in the held-out tissue split.

As in other recent work on DNA-based prediction tasks (5, 7, 9, 10) the sequence for each genomic site was obtained from human genome assembly hg19/GRCh37.

### Training the expression informed model

During training data was balanced per batch due to a 14:1 ratio of negative to positive examples. Each batch sampled an equal amount of accessible and non-accessible sites without replacement, such that one pass through all available negative training examples constituted multiple randomly permuted passes through all positive training examples. In situations where a DNase-seq file had more than a single matching RNA-seq file, sites from that DNase-seq file were randomly assigned to one of the multitude of corresponding RNA-seq expression vectors each time they were selected for a training batch.

To generate a validation set that was manageable to evaluate frequently we selected 40,000 random samples from each of accessible and non-accessible sites per validation DNase-seq file. This resulted in a set of 440,000 validation examples that were used to estimate ROC AUC throughout training.

However, upon stopping we also evaluated prediction performance across whole genomes (all potential DHSs) of all validation samples (Supplemental Table S2). In cases where multiple RNA-seq file matches existed, predictions across the entire genome were evaluated once for every possible DNase-seq and RNA-seq file pair. Whole genome evaluation gave a better characterization of performance on the intended application, especially as captured by PR AUC, which is less misleading in the presence of data imbalance. Results on the test sets were evaluated across whole genomes following the same procedure.

The total number of examples (all sites across all samples) for validation was 20.5 million and 22.2 million for testing in the held-out tissue partition.

All RNA-seq expression data used to train and test models was in units of log2(TPM + 1). There were many possible strategies for selecting the subset of genes for our input signature, but to initially avoid optimizing in this space, we relied on the prior work of the Library of Integrated Network-based Cellular Signatures (LINCS) and used their curated L1000 list of genes (49). To ensure that the models could be applied later to cancer genomes in TCGA we converted all L1000 gene names into Ensembl gene identifiers and kept only those genes that were available in both ENCODE and TCGA TOIL RNA-seq files. After this refinement, our final input L1000 gene list consisted of 978 genes.

### Expression informed model architecture and hyperparameters

We trained several alternative versions of our model and reported validation results over the course of training in Supplemental Figure S3 and Supplemental Figure S4.

The tissue-specific models demonstrated that multi-task outputs could share common convolutional layers and provide an accurate prediction of DNA accessibility across distinct sample types. Thus, we expected that if an input vector was discriminative of cell type it was likely to be sufficient to integrate it into the network after the convolutional layers. We evaluated adding a fully connected layer (depth = 500) before concatenating RNA-seq data to output from the convolutional layers, but found that it performed consistently worse (Supplemental Figure S3) than direct concatenation without the fully connected layer (Figure 1D).

Transfer learning consistently shortened the training time across model variants, and we found that using weights learned from the corresponding data partitions before final cleanup of revoked files was more effective on the validation set than was transfer of convolutional layer weights from the best tissue-specific model. However, our most impactful changes were increasing the batch size (from 128 to 512, and finally to 2048), and decreasing the learning rate (from 0.001 to 0.0001).

The tissue-specific models had multi-task outputs so that each training sample provided an information-rich gradient based on multiple labels for backpropagation. Since using RNA-seq inputs eliminated the need for multi-task outputs, each sample now only provided gradient feedback based on a single output. The batch size increase was intended to compensate for this change in output dimension to produce a more useful gradient for each batch.

The learning rate decrease, on the other hand, was guided by the observation that training was reaching a point of slow improvement before even a single full pass through all negative training examples. Our new dataset was also significantly larger than that used to train tissue-specific models.

We initialized our final expression-informed model (Figure 1D) with weights learned from the first iteration of the dataset, before erroneous revoked files were removed. In turn, those models were initialized with convolutional layer parameters from our best performing tissue-specific factorized convolutions model (Figure 1C). An effective batch size of 2048 was used for training (2 GPUs processing distinct batches of 1024), with an Adam (45) learning rate of 0.0001 and a 0.25 fraction of positive to negative samples in every batch.

### Expression informed model evaluation on ENCODE and genomic site annotations

ENCODE test set results were summarized in two ways: as a mean of AUC scores computed per whole genome sample (mean tissue type AUC in Tables 3 and 4), and as a single AUC score computed by considering predictions for all sites across all whole genome samples together (overall AUC in Tables 3 and 4). Only the latter was reported for performance analysis by genomic site type.

Two key sources were used to assign functional labels to accessibility prediction sites for performance breakdown. Exon, intragenic, and intergenic regions were derived from annotations defined by GENCODE v19 (50). Promoter and promoter flank, and enhancer region annotations were obtained from the Ensembl Regulatory Build (51).

When investigating correlation of training similarity to test sample performance, since the modulating factor between predictions applied to different tissues is the input RNA-seq data, distance between test samples, *t*, and the training set, *T*, was computed as d(*t*, *T*) = min_*i*∈*T*_‖***r***_*i*_ - ***r***_*t*_‖, where ***r***_*t*_ is a test sample’s vector of log_2_(TPM + 1) expression levels for all L1000 genes.

### Predicting DNA accessibility in TCGA

We applied our best expression informed model trained on the held-out tissue ENCODE partition to predict accessibility in TCGA. We restricted our predictions to promoter and promoter flank sites, since performance at those sites was high across all tests.

TOIL RNA-seq transcripts per million (TPM) gene expression data was used to obtain L1000 input gene signatures for all processed TCGA samples (32, 33). All expression values were converted from log_2_(TPM + 0.001) to log_2_(TPM + 1) before use.

For landscape views of accessibility and mutation impact analysis (Figure 3) we considered only samples with WGS available, and used mutation calls from an internal tool. For each sample site affected by at least one mutation, the change in predicted accessibility was computed before and after each mutation was applied, independently for SNPs and INDELs (Figure 3B). In order to apply t-SNE to generate the per-site landscape view (Figure 3D) we represented each site by a vector of binary accessibility decisions at that position across all selected TCGA samples with all mutations applied. All mutations were also applied when generating the per-patient t-SNE visualization (Figure 3F).

### Accessibility in LUAD

To assess the uniqueness in perspective of accessibility versus RNA-seq, all LUAD samples for which we had WGS data were clustered into two groups via K-means. For this, four data sources were used: accessibility predictions for promoter and promoter flank sites, TOIL log_2_(TPM + 1) RNA-seq gene expression data, HiSeqV2 log_2_(normalized count + 1) RNA-seq gene expression data (52) and TOIL log_2_(TPM + 1) RNA-seq gene expression data for all genes in the L1000 gene set used as inputs to our expression informed model. For the first three datatypes, we clustered samples based on the top 5% most variable sites (for accessibility) or genes (for TOIL and HiSeqV2) across the LUAD cohort, following the logic that the most highly variable sites may highlight the most dramatic differentially active pathways. For the L1000 genes, clustering was based on the entire set of gene expression levels. To show the difference quantitatively between cluster assignments across data types we used adjusted mutual information (1) (Figure 4B).

Exploration of pathway enrichment between the accessibility clusters was performed using Enrichr (31). Genes for enrichment analysis were selected by first eliminating all genes below a standard deviation threshold of 0.33 (in TOIL data) across the LUAD cohort (in HiSeqV2 data the equivalently selected standard deviation threshold was 1.0). This threshold was selected to include the main peak of gene standard deviation and exclude the peak around zero (Supplemental Figure S5), comprised of genes with little change or very low levels of expression. All remaining genes were then compared with a two-sided t-test between the two clusters and p-values were adjusted with Benjamini Hochberg (BH) correction. Due to the low number of WGS samples in either cluster (21 samples in C0 and 20 samples in C1) a more permissive false discovery rate of 0.25 was chosen as the cutoff for differential expression. In TOIL data, this procedure returned 512 genes upregulated in C0 and 857 genes upregulated in C1. In HiSeqV2 data, the same process yielded 344 upregulated genes in C0 and 339 in C1.

For comparison of tumor mutational burden (TMB) across clusters, TMB was computed as the total count of missense and nonsense mutations in each WGS sample.

When the patient analysis set was expanded to include all LUAD samples without WGS mutation information, clustering based on promoter and promoter flank accessibility predictions was repeated with the same procedure as before (Figure 4D).

To investigate whether the accessibility space appeared continuous along the dimensions of most variance across LUAD we used Principal Component Analysis (PCA) applied to all promoter and flank accessibility predictions to project each sample onto the first three principal components (Figure 4F).

### Correlating accessibility count with gene expression

Total accessibility count used to investigate gene correlations was computed as the total number of promoter and promoter flank sites predicted to be accessible after applying the binary decision threshold (at 80% precision) defined on ENCODE data. Again, only genes whose standard deviation was above 0.33 were considered for correlation analysis. Both Pearson and Spearman measures were evaluated, and the threshold for both measures was an absolute value above 0.4. All genes satisfying the threshold were analyzed for KEGG pathway enrichment with Enrichr (666 genes for Pearson correlation, and 418 genes for Spearman) (Supplemental Table S4 and S5)

### Accessibility analysis in immune cell driven clusters

LUAD samples were clustered into two groups using K-means on vectors of lymphoid (21 cell types) and myeloid (13 cell types) xCell estimates (39), revealing a survival difference (Supplemental Figure S6C). We noticed that a plane orthogonal to the first principal component (PC1) partitioned cluster labels when xCell vectors were reduced to three dimensions with PCA (Supplemental Figure S6A and B). To exclude cases of near ambiguous label assignment and focus on more prominent differences we removed samples within a small margin at the midpoint between clusters (in PC1). Margin size was equal to the standard deviation of the smallest cluster in the PC1 dimension (Supplemental Figure S6D and E). After ignoring margin samples, the survival difference of patients between clusters increased in significance (logrank test p = 6.7e-4) (Supplemental Figure S6F).
Total methylation for all LUAD samples was computed as the sum of values at all sites measured by the Infinium HumanMethylation450 BeadChip, available from TCGA (53). Total accessible site count considered all promoter and promoter flank sites, with binary class assignment based on the 80% precision threshold (Figure 5B).

For further analysis of accessibility, only sites previously determined as facultative were considered and all with low standard deviation (< 0.135) across LUAD (N = 512) were eliminated, to ignore cohort specific constitutive sites with some tolerance for noise. The threshold was selected so that at minimum 10 accessibility values at a site had to be distinct from the site’s values across the whole cohort (Supplemental Figure S7A).

Each accessibility prediction site was assigned to its nearest gene, according to distance in base pairs, as defined by GENCODE v19 (50). We considered only accessibility sites within 50,000 base pairs as having a valid correspondence to a gene (Supplemental Figure S7B). Significantly differentiated accessibility sites were then used to vote for candidate upregulated genes in each cluster. A very conservative significance threshold (two sided t-test BH adj. p < 1.0e-5) was selected so as to only focus on the most striking accessibility differences. Each site was allowed to contribute a single vote to its corresponding gene according to the cluster in which the site was more accessible.

Genes with a consistent direction of upregulation votes were considered cluster-specific candidate genes to test for differential expression (532 genes in X0 and 2250 genes in X1). From the candidate genes for each cluster that also had significant (two sided t-test BH adj. p < 0.01) differential expression (190 in X0 and 835 in X1) we identified the group in each cluster that was consistent (123 in X0, and 536 in X1) and inconsistent (67 in X0, and 299 in X1) with the direction of increased accessibility. All four sets were then tested for KEGG pathway enrichment via Enrichr (Supplemental Table S6 and S7).

### Predicting immune state from promoter and promoter flank accessibility

To train an ensemble of distinct models to discriminate immune hot from immune cold we used three fold cross validation; independently partitioning hot (X0 from immune cell based clustering) and cold (X1 from immune cell clustering) samples randomly to maintain an equal ratio across each fold. Training on different random subsets of data enhanced robustness when dealing with training label uncertainty. Each classifier was an RBF kernel SVM with *C* = 3.5, and 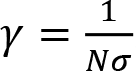, where *N* was the number of features and *σ* was the standard deviation of feature values across the training set. Additionally, training samples were balanced by weights inversely proportional to class frequency, and Platt scaling (54) was used to obtain probability estimates from SVM classification. During ensemble classifier application we excluded all samples that did not have a mean probability of at least 0.5 for the ensemble’s majority class prediction.

Input features to the classifier were binary accessibility predictions for a set of 484 sites comprised from the union of all immune hot (X0) sites consistent with gene expression and immune cold (X1) sites inconsistent with expression, as obtained from analysis of the xCell driven LUAD clusters. These sites were chosen both for their association with significant differences in expression of corresponding genes and the enrichment of those gene sets for immune relevant pathways.

Expanding the application domain of the immune activity classifier to previously unprocessed TCGA cohorts involved first applying our expression informed convolutional neural network model to all promoter and promoter flank sites in the new data. As previously, when expanding our LUAD sample size to non-WGS data, we used only the reference genome (hg19/GRCh37) and TOIL log_2_(TPM + 1) RNA-seq gene expression data for all predictions. Predictions that incorporated mutation information were included only for samples in our original six cohorts for which WGS was available.

## Supporting information

L1000 gene set ids

ENCODE accession ids for dataset splits

List of potential accessibility sites analyzed

ENCODE accession ids for dataset splits

List of potential accessibility sites analyzed

ENCODE accession ids for dataset splits

List of potential accessibility sites analyzed

ENCODE accession ids for dataset splits

List of potential accessibility sites analyzed

## Data Access

All raw data used in this study was obtained from public sources and all relevant accession numbers are made available as supplemental material.

