## Supplemental Figures and Tables for "Deep learning with implicit handling of tissue-specific phenomena predicts tumor DNA accessibility and immune activity"

### Dataset partitions

For the paired DNase-seq and RNA-seq dataset used to train our newly proposed model, we used two pairs of training and testing partitions. The first consisted of randomly held out samples at the DNase-seq file level (referred to as "tissue overlap" in the main text), while the second partitioning of the data specifically held out tissues for testing that were not included in training ("held-out tissue" throughout the main text). Both partition pairs are summarized by t-SNE plots in Figure S2 and the distribution of DNase-seq files for all tissues across partitions is summarized in Table S1.

Complete lists of all ENCODE file identifiers for each data partitioning used are provided as supplementary text files. This includes both generations of data partitions, before ("fold1v0_file_info.txt" and "fold2v0_file_info.txt") and after ("fold1v2_file_info.txt" and "fold2v2_file_info.txt") revoking of files by the ENCODE consortium. Lists of all potential accessibility sites used for training and prediction in each data partitioning scenario are also provided ("fold*_seq_sites.txt"). All files prefixed with "fold1" refer to the tissue overlap partitioning, while all files prefixed with "fold2" correspond to the held-out tissue set. Finally, we include an additional text file listing all genes used in the input RNA-seq signature ("ensembl_gene_ids_L1000.txt") derived from the Library of Integrated Network-based Cellular Signatures (LINCS) L1000 list of genes after conversion to Ensembl gene identifiers and checking for overlap with TCGA gene expression estimates from TOIL.

**Supplemental Figures**

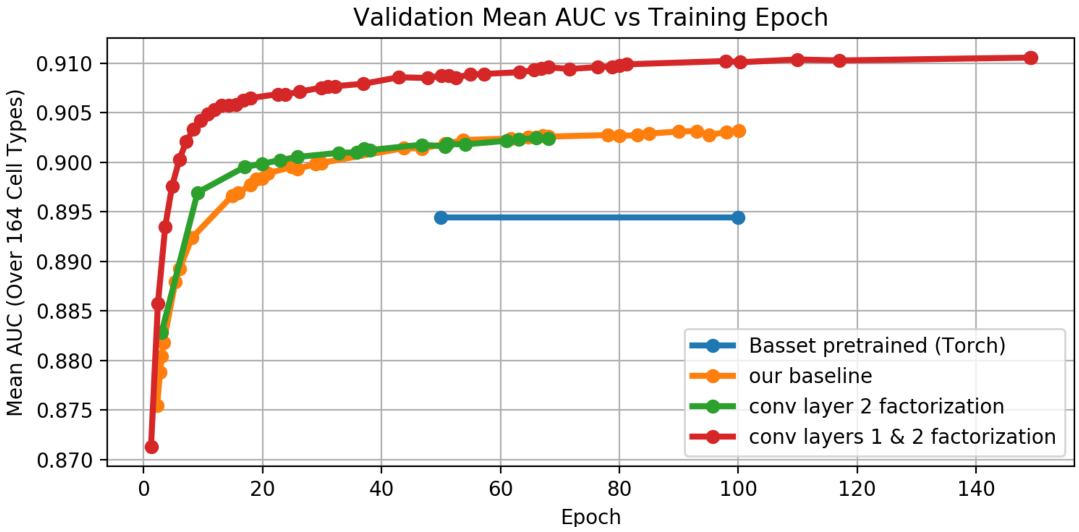


**Supplemental Figure S1. Training of tissue-type-specific model architectures**. The mean ROC AUC across 164 cell types in the validation set versus training epoch is shown. The result obtained by the pre-trained model provided by the authors of Basset is shown for reference, but since the number of training epochs was not reported, an arbitrary range was selected for display. We explored independent factorization of the second convolutional layer of the baseline model, and achieved the best performance when both the first and second convolutional layers were factorized.

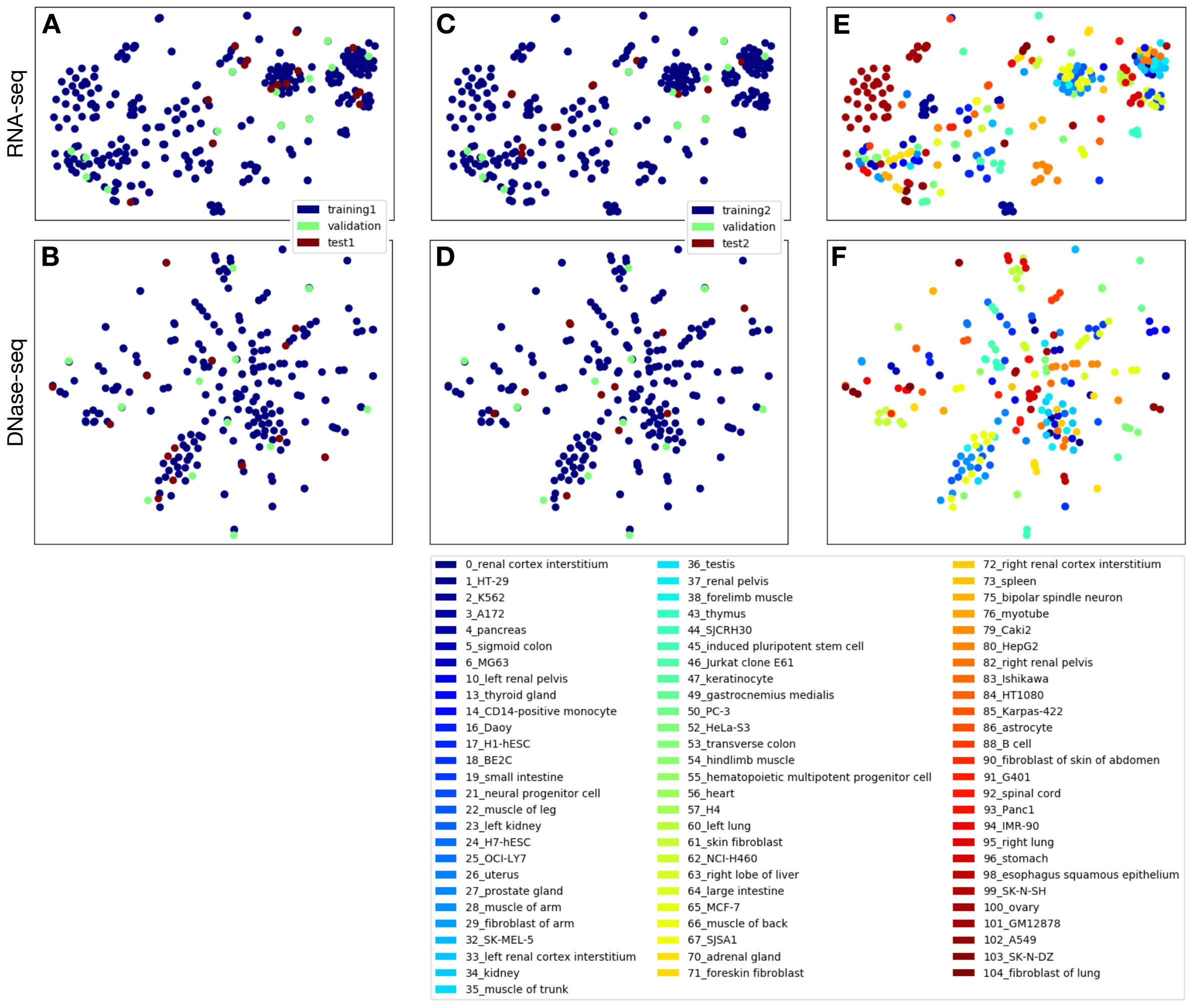


**Supplemental Figure S2. t-SNE embedding of ENCODE dataset partitioning in RNA-seq and DNase-seq space**. Sample distribution is illustrated by t-SNE embedding of the tissue overlap data partitions (training1 and test1) based on (*A*) RNA-seq $\log_2(\text{TPM} + 1)$ expression data and (*B*) DNase-seq peaks, as well as the held-out tissues data partitions (training2 and test2) based on (*C*) RNA-seq and (*D*) DNase-seq. The original ENCODE sample type labels are also illustrated for t-SNE embedded (*E*) RNA-seq and (*F*) DNase-seq samples, illustrating that samples of similar tissue or function often appear in proximity to each other across both data types.

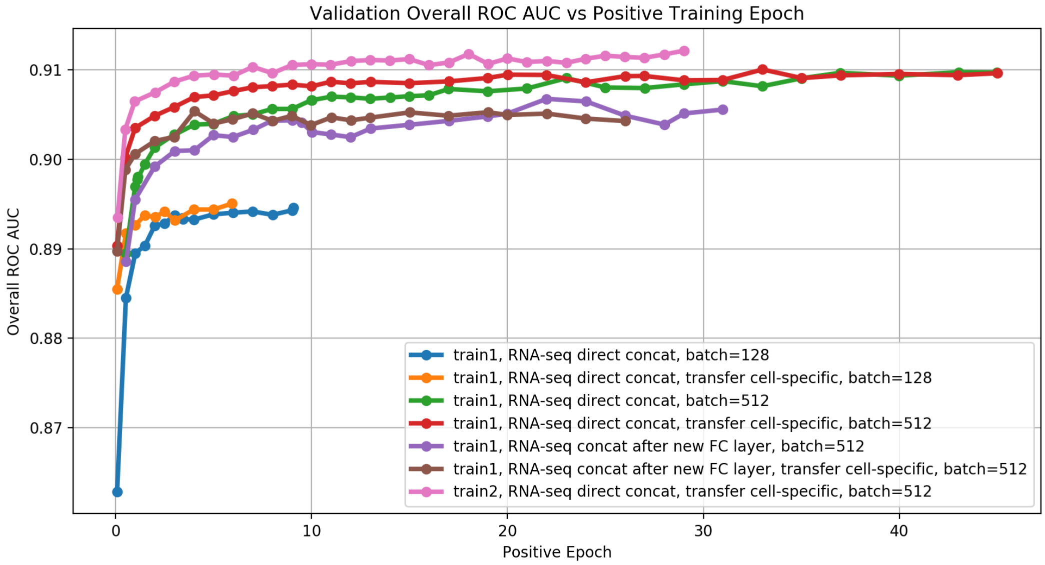


**Supplemental Figure S3. Overall ROC AUC for the small validation set**. The ROC AUC for the small validation set over number of passes through all positive examples (positive epochs) for several expression informed model architectures is shown. We experimented with adding a fully connected (FC) layer of depth 500 before concatenating (concat) gene expressions with outputs from the convolutional (conv) layers. However, increasing the batch size and initializing the convolutional layers with weights from our final tissue-specific model (transfer) improved performance most. Models trained on the tissue overlap set (train1) showed similar validation performance as those trained on the held-out tissue set (train2) with the same hyperparameters. This evaluation was done before the final dataset revision which revoked several suspected low quality samples, yet still provided valuable feedback for model selection.

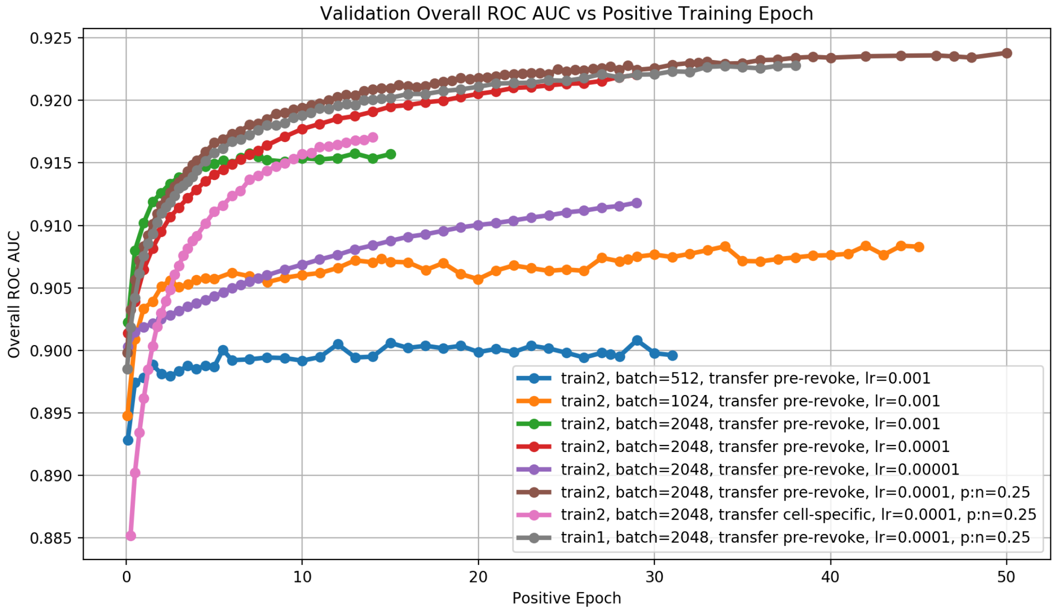


**Supplemental Figure S4. Overall ROC AUC for the small validation set over positive training epochs for models trained after the final dataset revision**. A further increase in batch size as well as a decreased learning rate (lr) led to additional significant improvements. Changing the fraction of positive samples per training batch (from p:n=0.5 to p:n=0.25) also slightly improved both ROC AUC as well as PR AUC in whole genome validation. Transfer of weights learned before final revoking of data (Figure S3) was a more effective initialization than weight transfer from our final tissue-specific model. Finally, we again confirmed that the same hyperparameters led to good validation performance across both training partitions: tissue overlap (train1) and held-out tissue (train2).

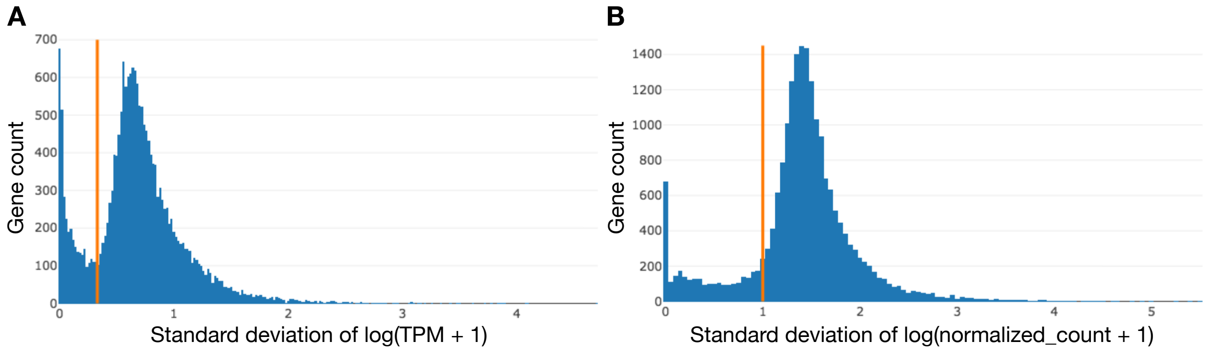


**Supplemental Figure S5, Histograms of gene level and expression SDs**. (A) Histogram of gene expression standard deviations across LUAD WGS samples in TOIL RNA-seq $log_2$(TPM + 1) data along with the selected threshold (0.33) in orange is shown. (B) Gene expression standard deviations across the same samples in HiSeqV2 $log_2$(normalized count+1) data and the selected threshold (1.0) in orange (B) are also shown. In both cases the thresholds eliminate genes with little change across samples or very low levels of expression, and keep all genes that constitute the main peak of values.

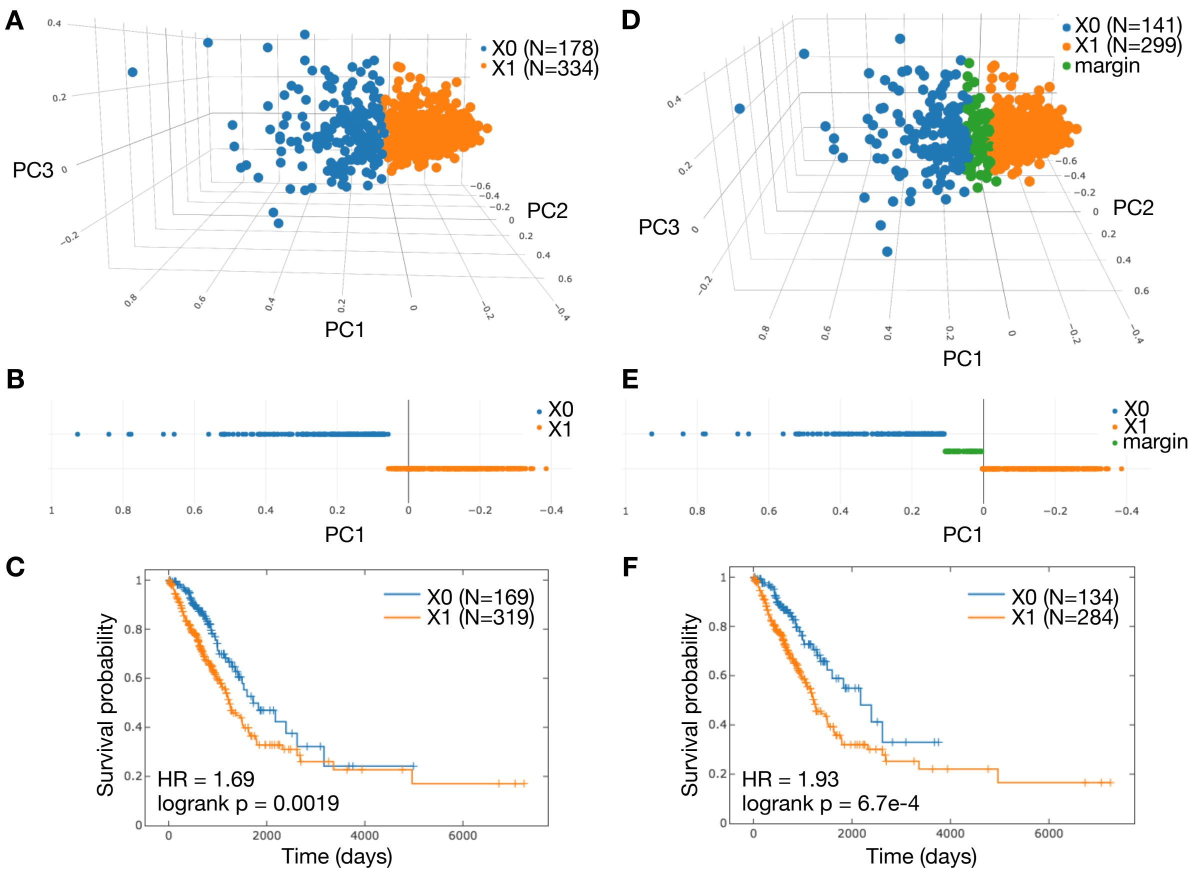


**Supplemental Figure S6. Adding a margin between immune cell based clusters in LUAD samples.** (A) All LUAD samples are shown with respect to the first three principal components (PC1-3) of their lymphoid and myeloid xCell estimates, colored according to labels assigned from k-means clustering. (B) Plotting the labeled data according to only the first principal component clearly shows the location of a separating plane between the clusters. The points excluded by introducing a margin at the scale of the smaller cluster's standard deviation along PC1 are shown (D) and (E). The impact on survival between X0 and X1 is shown by Kaplan-Meier plots before (C) and after (F) the margin was introduced. Kaplan-Meier plots are annotated with group size (N), logrank test p-values and hazard ratio (HR) based on a Cox proportional hazards (CoxPH) model regression using class assignment as the only explanatory variable.

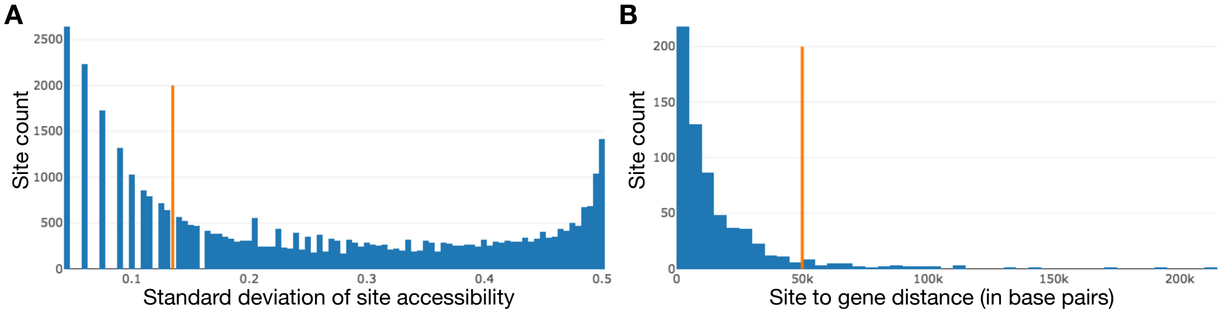


**Supplemental Figure S7. Histograms of accessibility SDs and site to gene distance.** (A) A histogram of the standard deviation of accessibility classifications in LUAD of promoter and promoter flank sites previously identified as facultative (40,823 sites) based on t-SNE across our initial set of six TCGA cohorts is shown. The threshold (< 0.135), in orange, identifies a subset of 25,093 sites facultative in LUAD. (B) The second histogram shows the site to nearest gene distances for all accessibility sites that also satisfy BH adjusted $p < 1.0e\text{-}5$ from a two-sided t-test between the LUAD immune cell driven clusters (3246 sites). Only sites within 50k base pairs (orange) were considered when voting for gene accessibility.

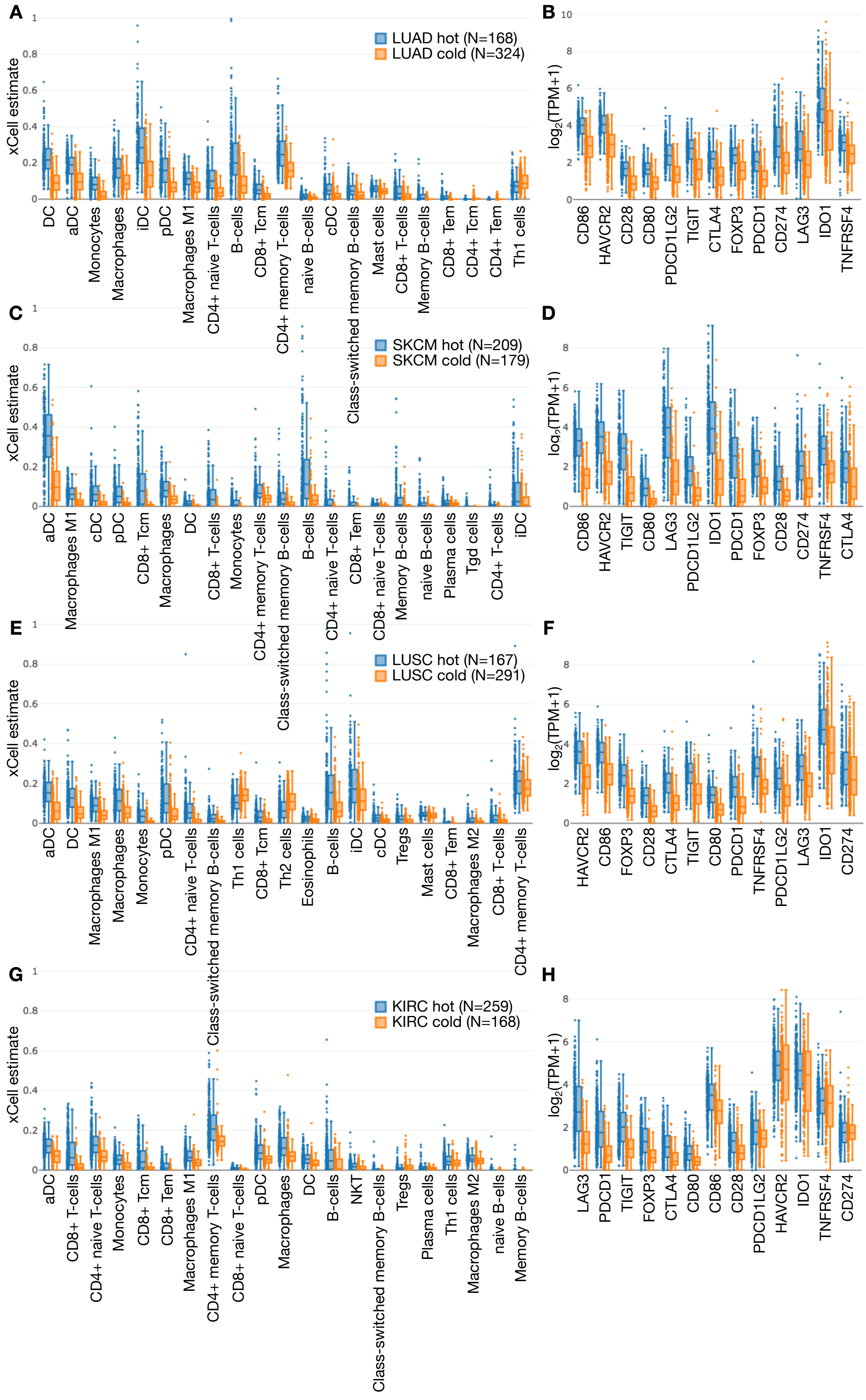

A
LUAD hot (N=168)
LUAD cold (N=324)
xCell estimate
DC
aDC
Monocytes
Macrophages
iDC
pDC
Macrophages M1
CD4+ naive T-cells
B-cells
CD8+ Tcm
CD4+ memory T-cells
naive B-cells
cDC
Class-switched memory B-cells
Mast cells
CD8+ T-cells
Memory B-cells
CD8+ Tem
CD4+ Tcm
CD4+ Tem
Th1 cells
B
log2(TPM+1)
CD86
HAVCR2
CD28
CD80
PDCD1LG2
TIGIT
CTLA4
FOXP3
PDCD1
CD274
LAG3
IDO1
TNFRSF4
C
SKCM hot (N=209)
SKCM cold (N=179)
xCell estimate
aDC
Macrophages M1
cDC
pDC
CD8+ Tcm
Macrophages
DC
CD8+ T-cells
Monocytes
CD4+ memory T-cells
Class-switched memory B-cells
B-cells
CD4+ naive T-cells
CD8+ Tem
CD8+ naive T-cells
Memory B-cells
naive B-cells
Plasma cells
Tgd cells
CD4+ T-cells
iDC
D
log2(TPM+1)
CD86
HAVCR2
TIGIT
CD80
LAG3
PDCD1LG2
IDO1
PDCD1
FOXP3
CD28
CD274
TNFRSF4
CTLA4
E
LUSC hot (N=167)
LUSC cold (N=291)
xCell estimate
aDC
DC
Macrophages M1
Macrophages
Monocytes
pDC
CD4+ naive T-cells
Class-switched memory B-cells
Th1 cells
CD8+ Tcm
Th2 cells
Eosinophils
B-cells
iDC
cDC
Tregs
Mast cells
CD8+ Tem
Macrophages M2
CD8+ T-cells
CD4+ memory T-cells
F
log2(TPM+1)
HAVCR2
CD86
FOXP3
CD28
CTLA4
TIGIT
CD80
PDCD1
TNFRSF4
PDCD1LG2
LAG3
IDO1
CD274
G
KIRC hot (N=259)
KIRC cold (N=168)
xCell estimate
aDC
CD8+ T-cells
CD4+ naive T-cells
Monocytes
CD8+ Tcm
CD8+ Tem
Macrophages M1
CD4+ memory T-cells
CD8+ naive T-cells
pDC
Macrophages
DC
B-cells
NKT
Class-switched memory B-cells
Tregs
Plasma cells
Th1 cells
Macrophages M2
naive B-ells
Memory B-cells
H
log2(TPM+1)
LAG3
PDCD1
TIGIT
FOXP3
CTLA4
CD80
CD86
CD28
PDCD1LG2
HAVCR2
IDO1
TNFRSF4
CD274

**Supplemental Figure S8. Example immune cell and checkpoint gene distributions across predicted hot and cold tumors.** Examples of xCell estimates and checkpoint gene expression levels compared across multiple cohorts for tumors classified by our 3 SVM ensemble as hot or cold, with group size shown (N). All x-axis labels are ordered by significance based on a two-sided t-test between tumors classified as hot and cold. Only the top 21 most significant xCell estimates from lymphoid and myeloid cell categories are shown. (A) The LUAD xCell estimate and (B) checkpoint gene distributions demonstrate how application of the classifier affected significance ordering from the raw training data illustrated in Figure 5. (C,D) SKCM provides an example of a distinct cohort in which immune hot patients exhibited longer survival. (E,F) LUSC demonstrates one case where immune activity appears to have no effect. (G,H) Finally, KIRC shows a case where immune active patients have significantly worse survival, and also exhibits a curious lack of difference between levels of CD274 (also called PD-L1) and PDCD1LG2 (also called PD-L2) compared to other cohorts split by our accessibility based immune activity classifier.

### Supplemental Tables

**Supplemental Table S1. Number of DNase-seq files by tissue name per dataset partition, including tissue overlap (TO) and held-out tissue (HOT) sets.**

| ENCODE Tissue Name | Validation | Train (TO) | Test (TO) | Train (HOT) | Test (HOT) |
|---|---|---|---|---|---|
| renal cortex interstitium | 0 | 3 | 0 | 3 | 0 |
| HT-29 | 0 | 2 | 0 | 2 | 0 |
| K562 | 0 | 2 | 0 | 2 | 0 |
| A172 | 0 | 1 | 1 | 2 | 0 |
| pancreas | 0 | 1 | 0 | 1 | 0 |
| MG63 | 0 | 2 | 0 | 2 | 0 |
| left renal pelvis | 0 | 3 | 1 | 4 | 0 |
| thyroid gland | 0 | 6 | 0 | 6 | 0 |
| CD14-positive monocyte | 1 | 1 | 0 | 1 | 0 |
| Daoy | 0 | 2 | 0 | 2 | 0 |
| H1-hESC | 0 | 2 | 0 | 2 | 0 |
| BE2C | 0 | 2 | 0 | 2 | 0 |
| small intestine | 0 | 3 | 2 | 5 | 0 |
| neural progenitor cell | 0 | 2 | 0 | 2 | 0 |
| muscle of leg | 0 | 7 | 0 | 7 | 0 |
| left kidney | 0 | 1 | 0 | 0 | 1 |
| H7-hESC | 0 | 3 | 0 | 3 | 0 |
| OCI-LY7 | 0 | 2 | 0 | 0 | 2 |
| uterus | 0 | 1 | 0 | 1 | 0 |
| prostate gland | 0 | 1 | 0 | 0 | 1 |
| muscle of arm | 2 | 7 | 1 | 8 | 0 |
| fibroblast of arm | 1 | 1 | 0 | 1 | 0 |
| SK-MEL-5 | 0 | 2 | 0 | 2 | 0 |
| left renal cortex interstitium | 0 | 4 | 0 | 4 | 0 |
| kidney | 0 | 4 | 0 | 4 | 0 |
| muscle of trunk | 0 | 1 | 0 | 1 | 0 |
| testis | 0 | 2 | 0 | 2 | 0 |
| renal pelvis | 0 | 3 | 0 | 3 | 0 |
| forelimb muscle | 0 | 0 | 1 | 1 | 0 |
| thymus | 0 | 6 | 0 | 6 | 0 |
| SJCRH30 | 1 | 2 | 0 | 2 | 0 |
| induced pluripotent stem cell | 0 | 2 | 0 | 2 | 0 |
| Jurkat clone E61 | 0 | 2 | 0 | 2 | 0 |
| keratinocyte | 0 | 1 | 1 | 2 | 0 |
| PC-3 | 0 | 2 | 0 | 2 | 0 |
| HeLa-S3 | 0 | 4 | 0 | 4 | 0 |
| hindlimb muscle | 0 | 1 | 0 | 0 | 1 |
| hematopoietic multipotent progenitor cell | 0 | 1 | 0 | 1 | 0 |
| heart | 0 | 2 | 0 | 2 | 0 |
| H4 | 0 | 2 | 0 | 2 | 0 |
| left lung | 0 | 6 | 0 | 6 | 0 |
| skin fibroblast | 0 | 8 | 1 | 9 | 0 |
| NCI-H460 | 0 | 2 | 0 | 2 | 0 |
| large intestine | 0 | 5 | 1 | 6 | 0 |
| MCF-7 | 0 | 4 | 0 | 4 | 0 |
| muscle of back | 0 | 8 | 2 | 10 | 0 |
| SJSA1 | 0 | 2 | 0 | 2 | 0 |
| adrenal gland | 0 | 5 | 1 | 6 | 0 |
| foreskin fibroblast | 1 | 1 | 0 | 1 | 0 |
| right renal cortex interstitium | 0 | 3 | 0 | 3 | 0 |
| spleen | 0 | 1 | 0 | 0 | 1 |
| bipolar spindle neuron | 0 | 2 | 0 | 2 | 0 |
| myotube | 0 | 2 | 0 | 2 | 0 |
| Caki2 | 0 | 2 | 0 | 2 | 0 |
| HepG2 | 0 | 4 | 0 | 4 | 0 |
| right renal pelvis | 1 | 3 | 0 | 3 | 0 |
| Ishikawa | 0 | 1 | 0 | 1 | 0 |
| HT1080 | 0 | 2 | 0 | 2 | 0 |
| Karpas-422 | 0 | 2 | 0 | 2 | 0 |
| astrocyte | 0 | 2 | 0 | 0 | 2 |
| B cell | 0 | 4 | 0 | 4 | 0 |
| fibroblast of skin of abdomen | 0 | 1 | 0 | 0 | 1 |
| G401 | 0 | 2 | 0 | 0 | 2 |
| spinal cord | 1 | 1 | 0 | 1 | 0 |
| Panc1 | 1 | 1 | 0 | 1 | 0 |
| IMR-90 | 0 | 4 | 0 | 4 | 0 |
| right lung | 1 | 3 | 0 | 3 | 0 |
| stomach | 0 | 5 | 0 | 5 | 0 |
| SK-N-SH | 0 | 2 | 0 | 2 | 0 |
| ovary | 0 | 1 | 0 | 1 | 0 |
| GM12878 | 0 | 2 | 0 | 2 | 0 |
| A549 | 1 | 2 | 0 | 2 | 0 |
| SK-N-DZ | 0 | 1 | 1 | 2 | 0 |
| fibroblast of lung | 0 | 5 | 1 | 6 | 0 |

**Supplemental Table S2. Whole genome validation results for our expression-informed model trained on tissue overlap (TO) and held-out tissue (HOT) sets.**

| Sample tissue type | ROC AUC (TO) | PR AUC (TO) | ROC AUC (HOT) | PR AUC (HOT) |
|---|---|---|---|---|
| CD14-positive monocyte | 0.888 | 0.559 | 0.889 | 0.563 |
| muscle of arm | 0.774, 0.959 | 0.654, 0.808 | 0.783, 0.960 | 0.671, 0.811 |
| fibroblast of arm | 0.898, 0.900 | 0.806, 0.809 | 0.898, 0.900 | 0.808, 0.811 |
| SJCRH30 | 0.875 | 0.727 | 0.875 | 0.730 |
| foreskin fibroblast | 0.953 | 0.774 | 0.953 | 0.771 |
| right renal pelvis | 0.967 | 0.833 | 0.968 | 0.836 |
| spinal cord | 0.947 | 0.714 | 0.946 | 0.713 |
| Panc1 | 0.957 | 0.713 | 0.958 | 0.711 |
| right lung | 0.958 | 0.781 | 0.958 | 0.782 |
| A549 | 0.902 | 0.735 | 0.900 | 0.734 |
| mean tissue type AUC | 0.915 | 0.743 | 0.916 | 0.745 |
| **overall AUC** | **0.912** | **0.721** | **0.913** | **0.725** |

**Supplemental Table S3. Enhancer results across held-out tissue test set whole genomes.**

| Sample tissue type | ROC AUC | PR AUC |
|---|---|---|
| left kidney | 0.934 | 0.737 |
| OCI-LY7 | 0.845, 0.845, 0.850, 0.850 | 0.645, 0.643, 0.606, 0.605 |
| prostate gland | 0.817 | 0.490 |
| hindlimb muscle | 0.933 | 0.908 |
| spleen | 0.809 | 0.471 |
| astrocyte | 0.931, 0.898 | 0.967, 0.833 |
| fibroblast of skin of abdomen | 0.953 | 0.940 |
| G401 | 0.640, 0.694 | 0.432, 0.270 |
| **overall AUC** | **0.870** | **0.732** |

**Supplemental Table S4. Pathway enrichment (Enrichr) results with adjusted p < 1.0e-4 for all 418 genes correlated with total promoter and promoter flank accessibility in LUAD with |Spearman correlation| > 0.4.**

| KEGG pathway | p | Adj. p | Z-score | Combined score |
|---|---|---|---|---|
| Osteoclast differentiation (hsa04380) | 3.74e-17 | 7.45e-15 | -1.86 | 70.51 |
| TNF signaling pathway (hsa04668) | 1.31e-10 | 1.30e-8 | -1.89 | 42.96 |
| Amoebiasis (hsa05146) | 2.48e-9 | 1.64e-7 | -1.82 | 36.03 |
| Pathways in cancer (hsa05200) | 2.33e-8 | 6.50e-7 | -1.95 | 34.27 |
| Tuberculosis (hsa05152) | 6.68e-9 | 2.66e-7 | -1.71 | 32.13 |
| Pertussis (hsa05133) | 4.72e-9 | 2.35e-7 | -1.64 | 31.46 |
| Regulation of actin cytoskeleton (hsa04810) | 2.61e-8 | 6.50e-7 | -1.75 | 30.52 |
| NF-kappa B signaling pathway (hsa04064) | 8.08e-9 | 2.68e-7 | -1.62 | 30.28 |
| Epstein-Barr virus infection (hsa05169) | 1.28e-6 | 2.83e-5 | -1.75 | 23.75 |
| PI3K-Akt signaling pathway (hsa04151) | 3.31e-6 | 5.21e-5 | -1.78 | 22.48 |
| Chemokine signaling pathway (hsa04062) | 2.11e-6 | 4.01e-5 | -1.69 | 22.15 |
| Influenza A (hsa05164) | 4.29e-6 | 6.10e-5 | -1.63 | 20.18 |
| Cytokine-cytokine receptor interaction (hsa04060) | 3.40e-6 | 5.21e-5 | -1.59 | 20.00 |
| AGE-RAGE signaling pathway in diabetic complications (hsa04933) | 8.58e-6 | 1.00e-4 | -1.60 | 18.72 |
| Jak-STAT signaling pathway (hsa04630) | 6.09e-6 | 7.58e-5 | -1.55 | 18.61 |
| Staphylococcus aureus infection (hsa05150) | 2.22e-6 | 4.01e-5 | -1.43 | 18.56 |
| Measles (hsa05162) | 5.70e-6 | 7.56e-5 | -1.50 | 18.17 |

**Supplemental Table S5. Pathway enrichment (Enrichr) results with adjusted p < 1.0e-6 for all 666 genes correlated with total promoter and promoter flank accessibility in LUAD with |Pearson correlation| > 0.4.**

| KEGG pathway | p | Adj. p | Z-score | Combined score |
|---|---|---|---|---|
| Osteoclast differentiation (hsa04380) | 4.35e-22 | 9.66e-20 | -1.86 | 91.69 |
| Influenza A (hsa05164) | 8.68e-13 | 9.63e-11 | -1.94 | 53.97 |
| TNF signaling pathway (hsa04668) | 1.37e-12 | 1.01e-10 | -1.86 | 50.83 |
| Regulation of actin cytoskeleton (hsa04810) | 5.54e-12 | 3.08e-10 | -1.85 | 47.86 |
| Hepatitis B (hsa05161) | 9.68e-11 | 4.30e-9 | -1.83 | 42.30 |
| HTLV-I infection (hsa05166) | 1.61e-10 | 5.95e-9 | -1.82 | 41.00 |
| Jak-STAT signaling pathway (hsa04630) | 5.20e-10 | 1.44e-8 | -1.75 | 37.47 |
| Pathways in cancer (hsa05200) | 1.76e-9 | 3.26e-8 | -1.82 | 36.61 |
| Chemokine signaling pathway (hsa04062) | 7.17e-10 | 1.77e-8 | -1.72 | 36.23 |
| Epstein-Barr virus infection (hsa05169) | 8.26e-10 | 1.83e-8 | -1.73 | 36.09 |
| Leukocyte transendothelial migration (hsa04670) | 3.16e-10 | 1.00e-8 | -1.57 | 34.40 |
| Tuberculosis (hsa05152) | 1.23e-9 | 2.48e-8 | -1.56 | 32.02 |
| Acute myeloid leukemia (hsa05221) | 3.69e-9 | 6.29e-8 | -1.55 | 30.04 |
| Measles (hsa05162) | 4.63e-9 | 7.15e-8 | -1.53 | 29.37 |
| Amoebiasis (hsa05146) | 4.83e-9 | 7.15e-8 | -1.48 | 28.35 |
| Viral carcinogenesis (hsa05203) | 2.30e-8 | 2.83e-7 | -1.55 | 27.26 |
| MAPK signaling pathway (hsa04010) | 3.37e-8 | 3.94e-7 | -1.54 | 26.49 |
| B cell receptor signaling pathway (hsa04662) | 1.42e-8 | 1.86e-7 | -1.46 | 26.36 |
| AGE-RAGE signaling pathway in diabetic complications (hsa04933) | 3.67e-8 | 4.07e-7 | -1.52 | 26.00 |
| NF-kappa B signaling pathway (hsa04064) | 1.01e-8 | 1.41e-7 | -1.34 | 24.68 |
| Focal adhesion (hsa04510) | 7.25e-8 | 7.31e-7 | -1.39 | 22.92 |
| Fc gamma R-mediated phagocytosis (hsa04666) | 6.71e-8 | 7.09e-7 | -1.29 | 21.35 |

**Supplemental Table S6. Top pathway enrichment (Enrichr) results for genes whose expression was consistent with increased accessibility in the immune active (C0) group of LUAD patients identified by xCell clustering.**

| KEGG pathway | p | Adj. p | Z-score | Combined score |
|---|---|---|---|---|
| Focal adhesion (hsa04510 | 0.000255 | 0.0355 | -1.92 | 15.90 |
| Osteoclast differentiation (hsa04380) | 0.00135 | 0.0936 | -1.84 | 12.15 |
| PI3K-Akt signaling pathway (hsa04151) | 0.00518 | 0.114 | -1.99 | 10.47 |
| Amoebiasis (hsa05146) | 0.00339 | 0.114 | -1.82 | 10.34 |
| Acute myeloid leukemia (hsa05221) | 0.00523 | 0.114 | -1.88 | 9.86 |
| Toxoplasmosis (hsa05145) | 0.00610 | 0.114 | -1.79 | 9.14 |
| Proteoglycans in cancer (hsa05205) | 0.00844 | 0.114 | -1.81 | 8.62 |
| Fc epsilon RI signaling pathway (hsa04664) | 0.00851 | 0.114 | -1.74 | 8.28 |
| Renal cell carcinoma (hsa05211) | 0.00784 | 0.114 | -1.67 | 8.09 |
| AMPK signaling pathway (hsa04152) | 0.00725 | 0.114 | -1.64 | 8.07 |
| Rap1 signaling pathway (hsa04015) | 0.00987 | 0.114 | -1.68 | 7.78 |
| Melanoma (hsa05218) | 0.00957 | 0.114 | -1.62 | 7.52 |

**Supplemental Table S7. Pathway enrichment (Enrichr) results (adj. p < 0.05) for genes whose expression was inconsistent with increased accessibility in the immune cold (C1) group of LUAD patients identified by xCell clustering.**

| KEGG pathway | p | Adj. p | Z-score | Combined score |
|---|---|---|---|---|
| Platelet activation (hsa04611) | 2.24e-6 | 4.38e-4 | -1.92 | 24.93 |
| Inflammatory mediator regulation of TRP channels (hsa04750) | 0.000112 | 0.0109 | -1.99 | 18.07 |
| Vascular smooth muscle contraction (hsa04270) | 0.000449 | 0.0235 | -1.82 | 14.00 |
| Chemokine signaling pathway (hsa04062) | 0.000529 | 0.0235 | -1.85 | 13.96 |
| cGMP-PKG signaling pathway (hsa04022) | 0.000944 | 0.0235 | -1.81 | 12.63 |
| Focal adhesion (hsa04510) | 0.000960 | 0.0235 | -1.77 | 12.31 |
| Intestinal immune network for IgA production (hsa04672) | 0.000731 | 0.0235 | -1.70 | 12.26 |
| Cholinergic synapse (hsa04725) | 0.00141 | 0.0247 | -1.83 | 12.04 |
| T cell receptor signaling pathway (hsa04660) | 0.000960 | 0.0235 | -1.69 | 11.72 |
| Calcium signaling pathway (hsa04020) | 0.00159 | 0.0247 | -1.68 | 10.83 |
| PI3K-Akt signaling pathway (hsa04151) | 0.00197 | 0.0263 | -1.73 | 10.78 |
| Phospholipase D signaling pathway (hsa04072) | 0.00148 | 0.0247 | -1.64 | 10.70 |
| Glutamatergic synapse (hsa04724) | 0.00164 | 0.0247 | -1.61 | 10.33 |
| Pathways in cancer (hsa05200) | 0.00275 | 0.0299 | -1.66 | 9.77 |
| ECM-receptor interaction (hsa04512) | 0.00144 | 0.0247 | -1.44 | 9.42 |
| Long-term depression (hsa04730) | 0.00202 | 0.0263 | -1.42 | 8.79 |
| Fc gamma R-mediated phagocytosis (hsa04666) | 0.00273 | 0.0299 | -1.41 | 8.30 |
| Rap1 signaling pathway (hsa04015) | 0.00461 | 0.0442 | -1.51 | 8.12 |
| Renin secretion (hsa04924) | 0.00268 | 0.0299 | -1.34 | 7.92 |
| B cell receptor signaling pathway (hsa04662) | 0.00474 | 0.0442 | -1.36 | 7.27 |
| Tuberculosis (hsa05152) | 0.00544 | 0.0484 | -1.29 | 6.74 |
| Leishmaniasis (hsa05140) | 0.00474 | 0.0442 | -1.21 | 6.49 |
